# Feeder-free culture of naive human pluripotent stem cells retaining embryonic, extraembryonic and blastoid generation potential

**DOI:** 10.1101/2025.01.17.633522

**Authors:** Giada Rossignoli, Michael Oberhuemer, Ida Sophie Brun, Irene Zorzan, Anna Osnato, Anne Wenzel, Emiel van Genderen, Andrea Drusin, Giorgia Panebianco, Nicolò Magri, Mairim Alexandra Solis, Chiara Colantuono, Sam Samuël Franciscus Allegonda van Knippenberg, Thi Xuan Ai Pham, Sherif Khodeer, Paolo Grumati, Davide Cacchiarelli, Paolo Martini, Nicolas Rivron, Vincent Pasque, Jan Jakub Żylicz, Martin Leeb, Graziano Martello

## Abstract

Conventional human pluripotent stem cells (hPSCs) are widely used to study early embryonic development, generate somatic cells, and model diseases, with differentiation potential aligned to a post-implantation epiblast identity. In the past decade, naive hPSCs, representing a pre-implantation stage, have been derived. Naive hPSCs efficiently differentiate towards embryonic and extraembryonic lineages such as trophectoderm, primitive endoderm, and extraembryonic mesoderm, and also self-organize into blastocyst-like structures called blastoids. However, their culture typically relies on mouse embryonic fibroblasts (MEFs), which are variable, resource-intensive, and can confound analyses. We report the long-term maintenance of naive hPSCs in a feeder-free, serum-coated system. We successfully expanded for up to 25 passages 8 different naive hPSCs lines across 5 laboratories. Growth rate, clonogenicity, and gene expression profiles on serum coating were comparable to MEF-based cultures, but serum coating eliminated fibroblast contamination. Naive hPSCs cultured on serum exhibited more efficient germ layer specification, retained trophectoderm potential and high blastoid formation efficiency. Exome sequencing revealed fewer mutations in serum-cultured cells, and mass spectrometry identified extracellular matrix proteins such as vitronectin, fibronectin, and collagens in the serum coating. Overall, serum coating offers a scalable, cost-effective and therefore widely applicable alternative for naive hPSC culture, maintaining developmental potential, reducing DNA mutations, and eliminating MEF-related confounding factors. We believe serum coating will expand the use of naive hPSCs to large-scale studies and facilitate the investigation of mechanistic insights into developmental and disease modelling.

## Introduction

The derivation of human pluripotent stem cells (hPSCs) has transformed the stem cell field. Given their potential to self-renew and differentiate into specialised cell types, there have been high expectations for their use in biological and medical research. hPSCs were first derived from the inner cell mass (ICM) of *in vitro* fertilised embryos (Human Embryonic Stem Cells, hESCs), and cultured in serum-based media on a feeder layer of inactivated mouse embryonic fibroblasts (MEFs)^1^, following the methods developed for ICM-derived mouse ESCs (mESCs)^2^. However, it soon became clear that the self-renewal requirements of hESCs and mESCs were different. Undifferentiated mESCs could be maintained in feeder-free conditions in the presence of leukaemia inhibitory factor (LIF) and foetal bovine serum (FBS) on gelatin-coated plates^3,4^. Subsequently, two inhibitors (2i), the MEK inhibitor PD0325901 and the GSK3 inhibitor CHIR99021 were shown to promote efficient mESCs self-renewal in the absence of feeders^5,6^. The combination of 2i and the cytokine LIF (2iL) resulted in more robust proliferation, increased oxidative phosphorylation and genome hypomethylation^7–9^. Since 2007, when hPSCs were reprogrammed from human somatic cells (human induced pluripotent stem cells, hiPSCs) by overexpressing a cocktail of transcription factors (OCT4, SOX2, KLF4 and c-MYC from Takahashi *et al*^10^, and OCT4, SOX2, NANOG and LIN28 from Yu *et al*^11^), hundreds of lines of hPSCs have been derived. Surprisingly, hPSCs were found not responsive to 2iL^12^. In contrast, they can be cultured without feeders on Matrigel or vitronectin-coated plates in Essential 8 (E8) or mTeSR media, both based on FGF2 and TGFβ supplementation^13–15^.

Although hESCs, like mESCs, are derived from pre-implantation blastocysts, they do not reside in a naive state of pluripotency and rather correspond to primed mouse ESCs derived from post-implantation epiblast (mEpiSCs), regarding growth factor dependence and transcriptional and epigenetic regulation^16–19^. Furthermore, hESCs (and hiPSCs) exhibit lower expression of pluripotency genes such as *NANOG*, *KLF4*, *KLF17*, *DPPA3*, and *DPPA5* when compared to the human naive epiblast^20^. The ability to convert mEpiSCs into mESCs provided a paradigm for the generation of naive human stem cells. This led different research groups to first derive naive hPSCs in 2iL by overexpressing transcription factors associated with naive pluripotency^12,21^, and later to implement more specific culture conditions and genetic manipulations^22–25^. Since then, naive hPSCs have been successfully derived from human blastocysts and somatic cell reprogramming by transient gene overexpression or chemical resetting from primed hPSCs in various media^26–30^. Successful maintenance of human naive pluripotency *in vitro* in a transgene-independent manner and in serum-free medium was initially achieved by culture optimisation starting from 2iL conditions^24,25^. The addition of Gö6983, a protein kinase C (PKC) inhibitor, previously shown to suppress also mESCs differentiation^31^, combined with a lower concentration of CHIR99021 (t2iLGö) was beneficial in maintaining compact colonies with morphology and proliferation of naive hESCs^24^. Later on, GSK3 inhibition was shown to be dispensable for the maintenance and resetting of naive hPSCs, so CHIR99021 was omitted and the tankyrase inhibitor XAV939 (PXGL) was added to achieve more robust and stable naive features^32^. A high-throughput chemical screen identified a combination of compounds, including the alternative GSK3 inhibitor IM12, the BRAF inhibitor SB590885, the SRC inhibitor WH-4-023, and the ROCK inhibitor Y-27632, supplemented with FGF and Activin A that synergise with PD0325901 and LIF to support the expansion of naive hPSCs (5i/L/AF)^25^. The addition of the JNK inhibitor SP600125 (6i/L/A) has been reported to increase the efficiency of naive hPSCs induction from the primed state^25^. Meanwhile, chemically defined conditions termed NHSM (naive human stem cell medium) were developed, in which 2iLGö was supplemented with p38 inhibitor SB203580 or BIRB796, JNK inhibitor SP600125, ROCK inhibitor Y-27632, bFGF and TGF-β1^23^.

The derivation of naive hPSCs has broadened the potential applications of human embryonic stem cells. Consistent with the fact that their transcriptomic state reflects a developmental state similar to the blastocyst stage epiblast rather than post-implantation epiblast, naive hPSCs exhibit a broader differentiation potential compared to primed hPSCs, enabling efficient differentiation into extraembryonic tissues, including trophectoderm (TE)^33–36^, primitive endoderm (PrE)^37,38^, and extraembryonic mesoderm^39^, three lineages previously inaccessible for *in vitro* studies. Furthermore, naive hPSCs can aggregate and generate 3D blastocyst-like structures composed of the three founding lineages as a model to study human implantation and peri-implantation development^40–42^. Naive hPSCs are characterised by global genomic hypomethylation compared to primed cells^43^ - like mESCs^9^ - and biallelic expression of multiple imprinted genes^44,45^. Naive hPSCs also use a bivalent metabolic system with a greater reliance on oxidative metabolism, whereas primed hPSCs are almost exclusively glycolytic^24,25,46,47^, as reported for naive mESCs^8^.

A major hurdle for studying the human naive state is that, unlike the mouse naive and human primed states, they are still routinely cultured on MEFs. In addition to the experimental cost and time involved, the quality of culture on feeders has been shown to depend on numerous other factors such as embryo age, passage number of MEFs and their genetic background, all of which affect gene expression and factor release from MEFs themselves, as well as stem cells proliferation and colony formation^48–51^. The exchange of signals between feeders and PSCs, some of which are still unknown, undoubtedly prevents a stringent control of the system and makes it more difficult to understand the best conditions and underlying biological pathways necessary for maintaining pluripotency *in vitro*, with implications also in downstream analyses and applications of these cells. There have been extensive efforts to eliminate feeders from naive hPSC cultures. However, to this date, they have not been widely adopted for multiple reasons. A 3D culture system based on Matrigel allows for a robust expansion of naive hPSCs in PXGL^52^. While it allows for efficient 3D differentiation, it is time- and resource-intensive as a routine culture method. Reduced proliferation was reported if MEFs were replaced with Matrigel or laminin-511 coating in t2iLGö medium in 2D^24^. Media formulations with extensive addition of inhibitors allowed for robust expansion of hPSCs on Matrigel or Vitronectin-coated plates (FINE, NHSM and RSeT, a commercial medium based on NHSM)^23,53^. However, these inhibitors affect the morphology^53^ or stabilise a transcriptional state of hPSCs distinct from naive cells routinely used in blastoid formation^23,27^. Consequently, the routinely used conditions for naive hPSCs expansion and blastoid generation rely on MEFs^40–42^.

The addition of serum to cell culture media has been used to improve cell adhesion since the first half of the last century^54–56^. Analysis of serum composition led to the discovery of the extracellular matrix (ECM) protein vitronectin^57^, together with fibronectin^58^. Murine EpiSCs can be expanded efficiently on FBS-coated plates^16^. Interestingly, mESCs cultured on serum-coated substrates in the presence of LIF maintain the expression of *OCT4* and grow as round cells in small colonies^59^. Moreover, while mESCs strongly attach when converted from 2iL to serum/LIF, cells in a serum-containing medium might have attachment problems when converted to serum-free 2iL, and the addition of a small amount of FBS in 2iL generally enhances adhesion^60^. Furthermore, conventional hPSCs have been successfully grown on serum-coated substrates without feeders^61^.

In this study, we observed, across 5 laboratories, that naive hPSCs of both induced and embryonic origins can be easily adapted to a feeder-free, serum-based coating in PXGL. Cells grown on serum coated culture dishes retain a naive pluripotency gene expression signature, and a proliferation rate and clonogenic capacity similar to the original lines on MEFs, without acquiring pathogenic-associated mutations. In addition, naive hPSCs cultured on serum coating retain full differentiation potential towards both extra-embryonic and embryonic tissues as well as the capacity to self-assemble into blastoids.

## Results

### Naive hPSCs spontaneously adapt to serum coating

Naive hiPSCs directly reprogrammed from somatic cells^30^ (HPD06 and HPD03) were plated in PXGL medium on dishes coated with 10% FBS diluted in DMEM (i.e. serum coating) without MEFs. Feeder-free converted cells retained their distinctive dome-shaped morphology (Supplementary Figure 1A) and expressed general pluripotency (*POU5F1/OCT4* and *NANOG*) and naive-specific markers (*TFCP2L1*, *KLF4* and *KLF17*) at comparable or even higher levels than cells cultured on MEFs, as well as low or undetectable expression of primed genes *OTX2* and *ZIC2* (Figure 1A). Comparable protein levels of naive markers KLF17 and TFCP2L1, and the general pluripotency markers OCT4 and NANOG were observed in feeder-free and MEFs cultured cells (Figure 1B, left, and Supplementary Figure 1B). Two naive PSC lines (HPD06 and HPD03) displayed a decrease in the proliferation rate (Supplementary Figure 1D) and clonogenic capability (Supplementary Figure 1E) when plated at low density on serum coating in comparison to the same cell lines on MEFs during the first 4 passages of the conversion. Upon long-term culture in feeder-free conditions, these hPSC lines retained the dome-shaped morphology (Figure 1C, left) and recovered growth rate and clonogenicity to levels comparable to the same cell line cultured on MEFs (Figure 1D-E). We conclude that serum coating allows for the maintenance of naive pluripotency markers and the long-term expansion of naive hiPSCs.

**FIGURE 1.**
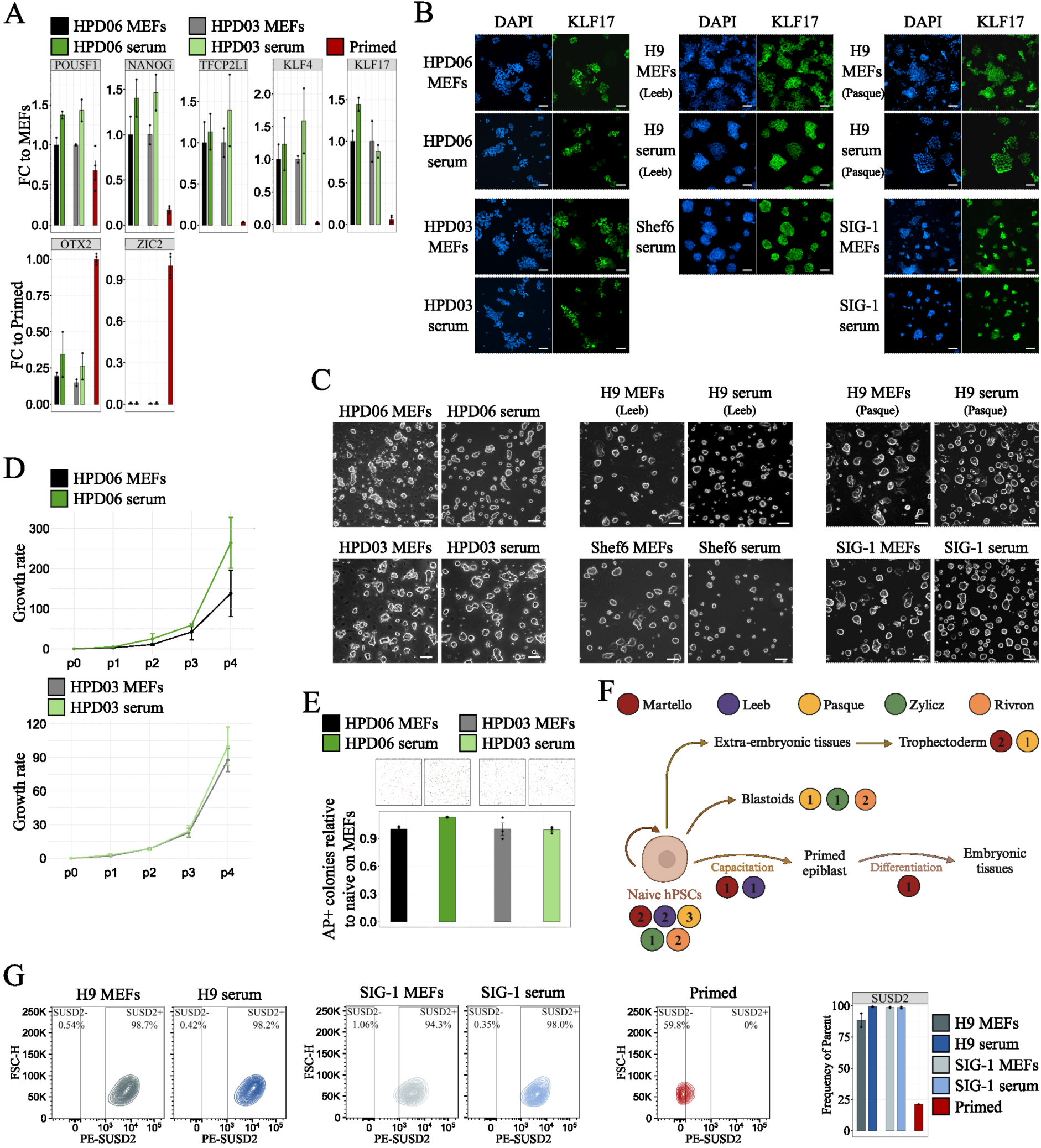
Serum coating allows for the maintenance of naive hPSCs. **A** Gene expression analysis by RT-qPCR of general (*POU5F1* and *NANOG*), naive (*TFCP2L1*, *KLF4* and *KLF17*) and primed (*OTX2* and *ZIC2*) pluripotency markers in HPD06 and HPD03 naive hiPSCs stably cultured on MEFs, serum or matrigel coating. Bars indicate the mean ± SEM of biological replicates shown as dots. n = 2 from independent experiments for naive hiPSCs and n = 4 from independent experiments for primed hPSCs. **B** Immunostaining for KLF17 of different naive hPSC lines (HPD06, HPD03 and SIG-1 iPSCs, and H9 and Shef6 ESCs) stably cultured on MEFs or serum coating. See also Supplementary Figure 1B-C. Scale bars: 100 μM. Representative images of two independent experiments are shown. **C** Morphologies of different naive hPSC lines (HPD06, HPD03 and SIG-1 iPSCs, and H9 and Shef6 ESCs) stably cultured on MEFs or serum coating. Scale bars: 100 μm. Representative images of two independent experiments are shown. **D** Growth rate of HPD06 and HPD03 naive hiPSCs stably cultured on MEFs or serum coating. Bars indicate the mean ± SEM of n = 2 independent experiments shown as dots. **E** Top: Representative AP staining images after clonal assay of HPD06 and HPD03 naive hiPSCs stably cultured on MEFs or serum coating. Bottom: Quantification of the relative number of AP-positive pluripotent colonies counted per well. Bars indicate the mean ± SEM of n = 3 independent experiments shown as dots. **F** Schematic representation of the analyses performed in this study. Coloured dots identify the laboratories involved, with numbers indicating cell lines used for each analysis. **G** Flow cytometry analysis of naive H9 hESCs and SIG-1 hiPSCs stably cultured on MEFs or serum coating. Left: Representative contour plots of SUSD-PE *versus* forward scatter. Right: Quantification of SUSD2 positive cells as the frequency of live cells. Bars indicate the mean ± SEM of n = 2 technical replicates shown as dots.

Prompted by these results, we decided to test extensively whether serum coating could be used for naive PSC lines of different origin (i.e. reprogramming of fibroblasts, resetting of conventional PSCs and embryo-derived), of both sexes and in different laboratories. Furthermore, we reasoned that an extensive test of the differentiation potential should be performed, including both embryonic and extraembryonic lineages, to test if serum coating allows for the maintenance of *bona fide* naive PSCs (Figure 1F). We successfully converted 8 human naive PSC lines in 5 different laboratories to feeder-free conditions (Table 1). They all displayed comparable expression of nuclear (OCT4 and NANOG) and naive markers (KLF17 and SUSD2) (Figure 1B, G and Supplementary Figure 1C, F), with dome-shaped morphology and robust expansion (Figure 1C). Although only HPD06 and HPD03 lines exhibited a lag phase before complete adaptation, we have not experienced a line failing to convert to serum coating in PXGL.

**TABLE 1.** Naive hPSC lines used in this study.

| Cell line | Origin | Sex | Derivation | Group |
| --- | --- | --- | --- | --- |
| HPD06 | Induced from human fibroblasts | Male | Somatic cell reprogramming<sup>30</sup> | Martello |
| HPD03 | Induced from human fibroblasts | Male | Somatic cell reprogramming<sup>30</sup> | Martello |
| H9<sup>1</sup> | Embryonic | Female | Epigenetic resetting<sup>106</sup> | Leeb |
| Shef6<sup>85</sup> | Embryonic | Female | Epigenetic resetting<sup>106</sup> | Leeb |
| H9<sup>1</sup> | Embryonic | Female | Epigenetic resetting<sup>106</sup> | Pasque |
| SIG-1 | Induced from human fibroblasts | Female | mRNA KLF4 reprogramming<sup>27</sup> | Pasque |
| HNES1 | Embryonic | Male | From blastocyst<sup>26</sup> | Pasque |
| H9<sup>1</sup> | Embryonic | Female | Epigenetic resetting<sup>106</sup> | Zylicz |
| KOLF2.1J | Induced from human fibroblasts | Male | Episomal Sox2-17 and KLF4 reprogramming<sup>107</sup> | Rivron |
| SCTi003-A | Induced from human fibroblasts | Female | Episomal Sox2-17 and KLF4 reprogramming<sup>107</sup> | Rivron |

We concluded that naive hPSCs spontaneously adapt to the feeder-free culture on serum coating in PXGL without additional treatment or genetic manipulation.

### Proteomics analysis of serum composition reveals abundant ECM proteins

We wondered how serum coating could sustain the attachment and growth of naive hPSCs. Previous studies identified Vitronectin as a component of serum accounting for its capacity to promote cell adhesion^57^. Therefore we assessed the composition of the coating through a label-free quantitative proteomics analysis. The entire list of identified proteins was screened and annotated for matrisome and matrisome-associated macromolecules using the Matrisome AnalyseR tool from The Matrisome Project^62^. Of the 850 proteins detected in serum coating, 183 (more than 20%) were classified as “Core matrisome” or “Matrisome-associated” hits. ECM glycoproteins and ECM-affiliated proteins and regulators accounted for the majority of the hits in these two categories (Figure 2A). The analysis of the top 100 most prevalent proteins in serum coating showed that Fibronectin, Vitronectin, Laminin, and several Collagens were present (Figure 2B).

**FIGURE 2.**
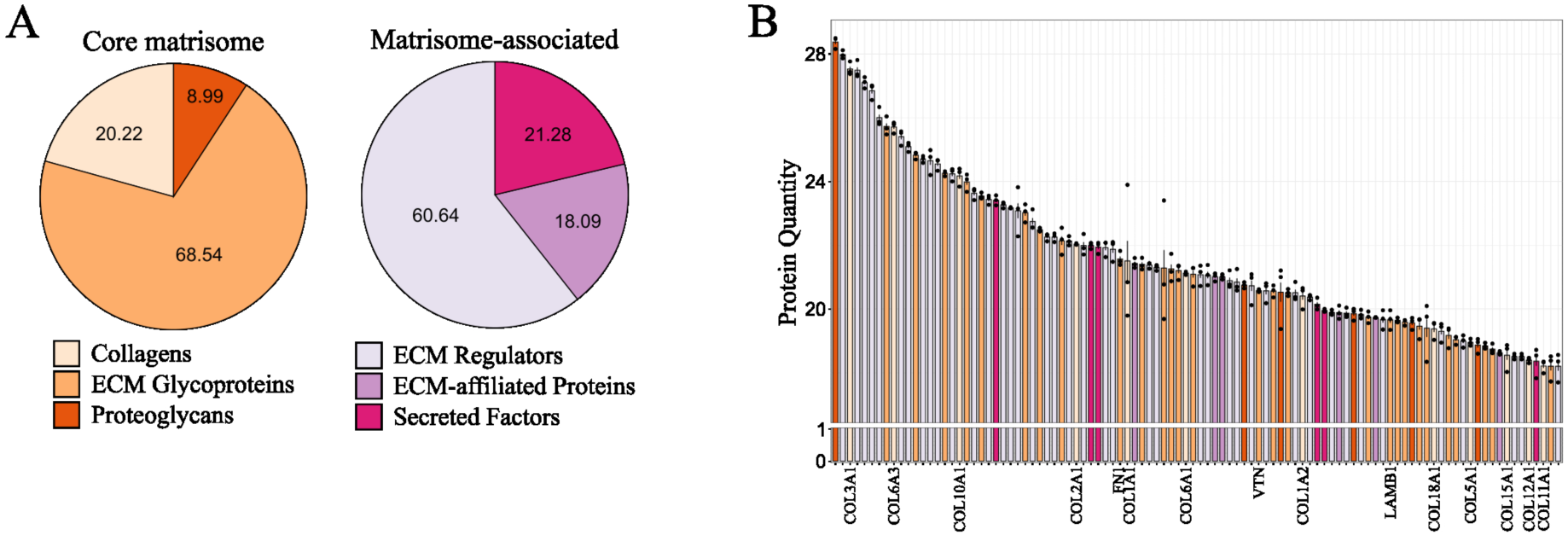
Serum contains a mixture of ECM proteins. **A** ECM proteins enrichment analysis of the serum coating proteome. Left: Composition of core matrisome proteins expressed in percentage. Right: Composition of matrisome-associated proteins expressed in percentage. **B** Top 100 most abundant proteins of the serum coating. Commonly used surface coatings for feeder-free cultures and detected collagens are highlighted. Bars indicate the mean ± SEM of n = 3 technical replicates shown as dots.

Therefore, the serum contains a mixture of ECM proteins previously reported to support conventional human PSCs^13–15^.

### Serum coating does not induce genetic variations in naive hPSCs

Several studies demonstrated that conventional hPSCs can acquire genetic changes during long-term *in vitro* culture, which commonly include mutations in cancer-associated genes, especially TP53^63,64^. We analysed specifically the two naive iPSC lines (HPD06 and HPD03) that required some passages to recover their proliferation rate on serum (Figure 1D), wondering whether they acquired mutations in hotspot genes during the adaptation. We collected genomic DNA from HPD06 and HPD03 cultured for more than 2 months after conversion to serum coating and from the same lines kept on MEFs (Figure 3A) and deep sequenced exomes (more than 600 million reads, 1000x independent base coverage). The comparison with the reference genome identified a total of 1649 shared variants (SNVs, indels, and CNVs) in protein-coding regions between HPD06 naive hPSCs cultured on MEFs and serum, likely accumulated during derivation and previous culture on MEFs (Figure 3B, left Venn diagrams). However, HPD06 continuously cultured on MEFs accumulated 936 variants, contrary to the 356 variants specifically detected in the same line cultured on serum. Similarly, more variants were detected in HPD03 naive hPSCs on MEFs than in the same line converted to serum coating (509 and 311, respectively; Figure 3B, top Venn diagrams). We then grouped protein-coding variants according to their clinical impact (tier I, strong significance; tier II, potential clinical significance; tier III, unknown clinical significance; and tier IV, benign or likely benign)^65^. For both HPD06 and HPD03, no Tier I alterations were identified and Tier IV represented the vast majority of detected variants (Figure 3B, bottom bar plots). Interestingly, for both naive lines, the culture on MEFs showed an accumulation of Tier III variants compared to serum coating (Figure 3B, bar plots). Few cancer-related genes previously reported as commonly mutated in conventional hPSCs showed only Tier IV benign or likely benign alterations (*FAT1*, *ASXL1* and *NF1*)^64^. No genetic alteration was found for *TP53*, a predominantly mutated gene during *in vitro* culture^63,64^.

**FIGURE 3.**
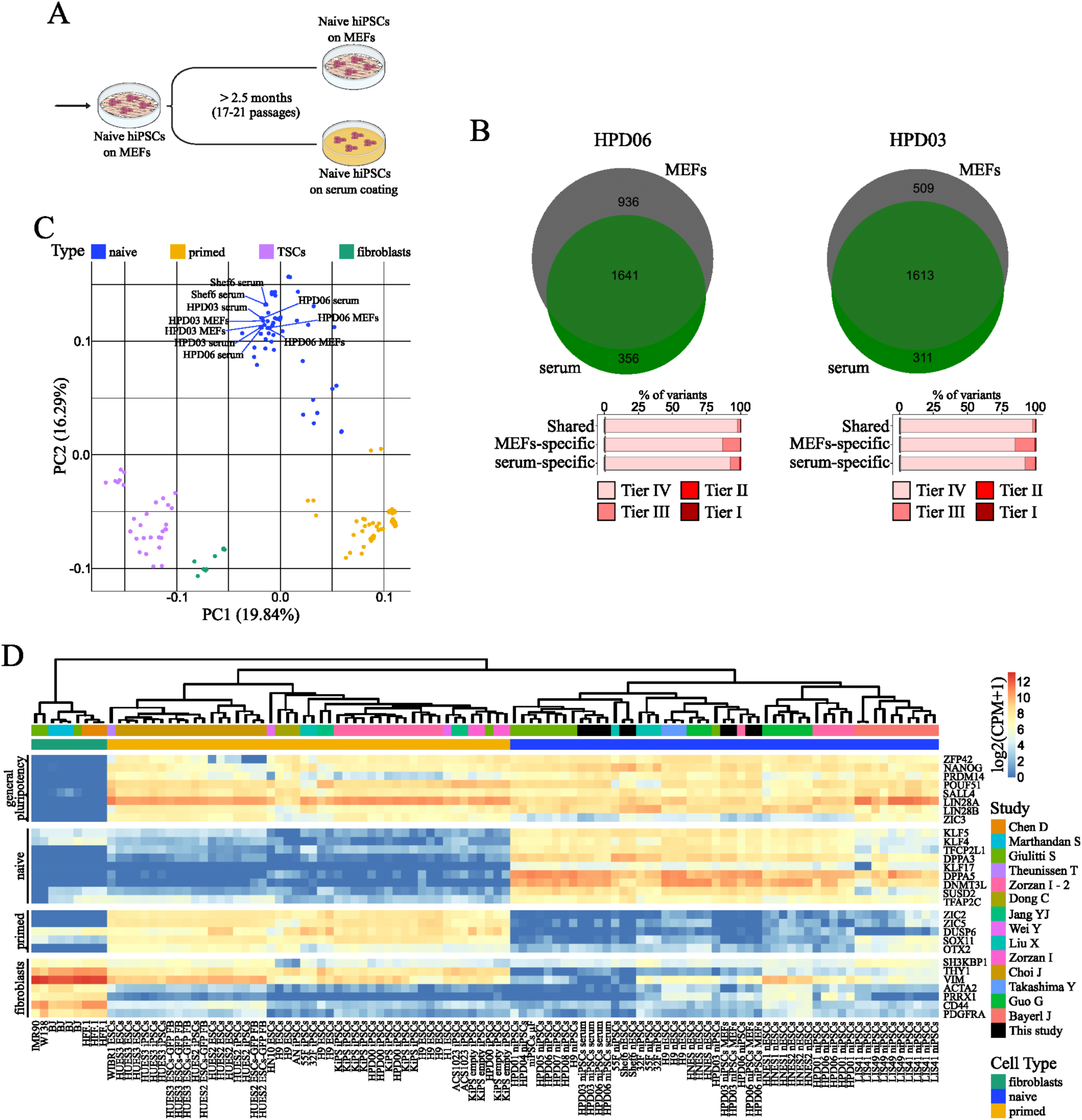
Naive hPSCs on serum coating do not accumulate pathogenic mutations and maintain their transcriptomic profile. **A** Schematic representation of the experimental setting for the Exome-Seq analysis of HPD06 and HPD03 naive hiPSCs stably cultured on MEFs or serum coating. **B** Analysis of detected genetic variations in coding genes of HPD06 and HPD03 naive hiPSCs stably cultured on MEFs or serum coating compared to the reference human genome. Top: Venn diagrams showing the quantification of shared and culture-specific (MEFs or serum coating) detected variants and mutated genes identified in each condition. All Tier II detected genes and genes identified as commonly mutated in hPSCs^64^ are highlighted. Bottom: Barplots of the composition of shared and culture-specific (MEFs or serum coating) detected variants classified by Tier and expressed in percentage. **C** Principal component analysis of naive hPSC lines (HPD06 and HPD03 hiPSCs, and Shef6 hESCs) stably cultured on MEFs or serum coating with published naive and primed hPSCs, TSCs and fibroblast lines performed on the top 5,000 most variable genes identified through RNA-seq. **D** Heatmap of general pluripotency, naive, primed, and fibroblasts genes in naive hPSC lines (HPD06 and HPD03 hiPSCs, and Shef6 hESCs) stably cultured on MEFs or serum coating and in published fibroblasts, primed hPSCs and naive hPSCs.

Overall, culturing naive hPSCs on serum coating did not lead to pathogenic mutations. Moreover, mutation rates of naive lines cultured on serum were lower compared to MEFs-cultured cells.

### Naive hPSCs on serum coating maintain the transcriptomic profile and lack of gene imprinting

To further characterise the naive features of hPSCs on serum, we compared the transcriptome of our naive hPSCs cultured on MEFs and serum coating with a panel of published transcriptomes spanning human naive and primed PSCs, TSCs and fibroblasts. RNA-seq data confirmed a distinct global expression profile of our naive hPSC lines compared with primed hPSCs, TSCs and fibroblasts, as they fall together within the main naive cluster (Figure 3C). Gene expression analysis confirmed robust induction of several core and naive specific pluripotency genes (Figure 3D), and low to undetectable expression of primed-specific markers, in line with published transcriptomes. Generally, primed and naive hPSCs cultured on MEFs showed variable expression of fibroblast markers, possibly due to MEF contamination in collected cells (Figure 3D and Supplementary 2C). Fibroblast markers were barely detectable in naive lines cultured on serum.

It has been observed that mitochondrial activation and metabolic realignment occur concurrently with the generation and stabilisation of naive hPSCs^24^. Mouse ESCs use oxidative phosphorylation, while mEpiSCs and hPSCs are mostly glycolytic and have low ability for mitochondrial respiration^46^. We collected the list of oxidative phosphorylation genes from KEGG (ko00190 pathway; https://www.genome.jp/kegg/pathway.html) and added the complex IV cytochrome c oxidase (COX) genes. In naive hPSCs, we observed an increase in the expression of most genes involved in oxidative phosphorylation (Supplementary Figure 2A). Furthermore, in agreement with previously published data^24,46^, the expression of most of the COX genes was decreased in primed hPSCs compared to naive hPSCs, both on MEFs and serum coating.

Previous studies reported biallelic expression of some imprinted genes in human naive PSCs^44,45,66^. Therefore, we analysed our bulk RNA-Seq samples with BrewerIX^45^, running the analysis with the complete pipeline. We detected a significant biallelic expression of H19 and ZIM2/PEG3 specifically in all naive hiPSCs, either on serum or MEFs, but not in primed hiPSCs (Supplementary Figure 2B), as previously reported^44,67^. Overall, naive cells displayed similar allelic expression of imprinted genes on serum and MEFs.

We concluded that naive hPSCs cultured on serum coating maintain their distinctive transcriptome profile without significant alterations compared to naive hPSCs cultured on MEFs.

### Serum-adapted naive hPSCs faithfully recapitulate embryonic differentiation trajectories

Upon withdrawal of self-renewal cues, naive hESCs exit the naive pluripotent state. Before gaining the ability to commit to embryonic cell fates, they need to undergo a process termed capacitation^68^. Capacitation requires approximately eight to ten days, consistent with the timing between the blastocyst and the pre-gastrulation post-implantation embryo *in vivo*. Hence, to assess the capacitation and differentiation potential of serum-cultured naive hPSCs (Figure 1F), we subjected them to a variety of differentiation conditions. Changes in morphology were equivalent between naive hPSCs cultured on MEFs or serum coating, across multiple differentiation media (Supplementary Figure 3A). Furthermore, expression of naive specific markers (*TFCP2L1*, *KLF4* and *NANOG*) was downregulated while expression of primed markers (*SALL2*, *CDH2* and *ZIC2)* was induced during the exit from naive pluripotency irrespective of whether hESCs were maintained on MEFs or serum before the induction of differentiation (Figure 4A). Consistently, naive transcription factors downregulation and primed marker upregulation were also observed in Shef6 hESCs cultured on serum coating inducing capacitation (Figure 4B) and were confirmed at the protein level (Supplementary Figure 3B).

**FIGURE 4.**
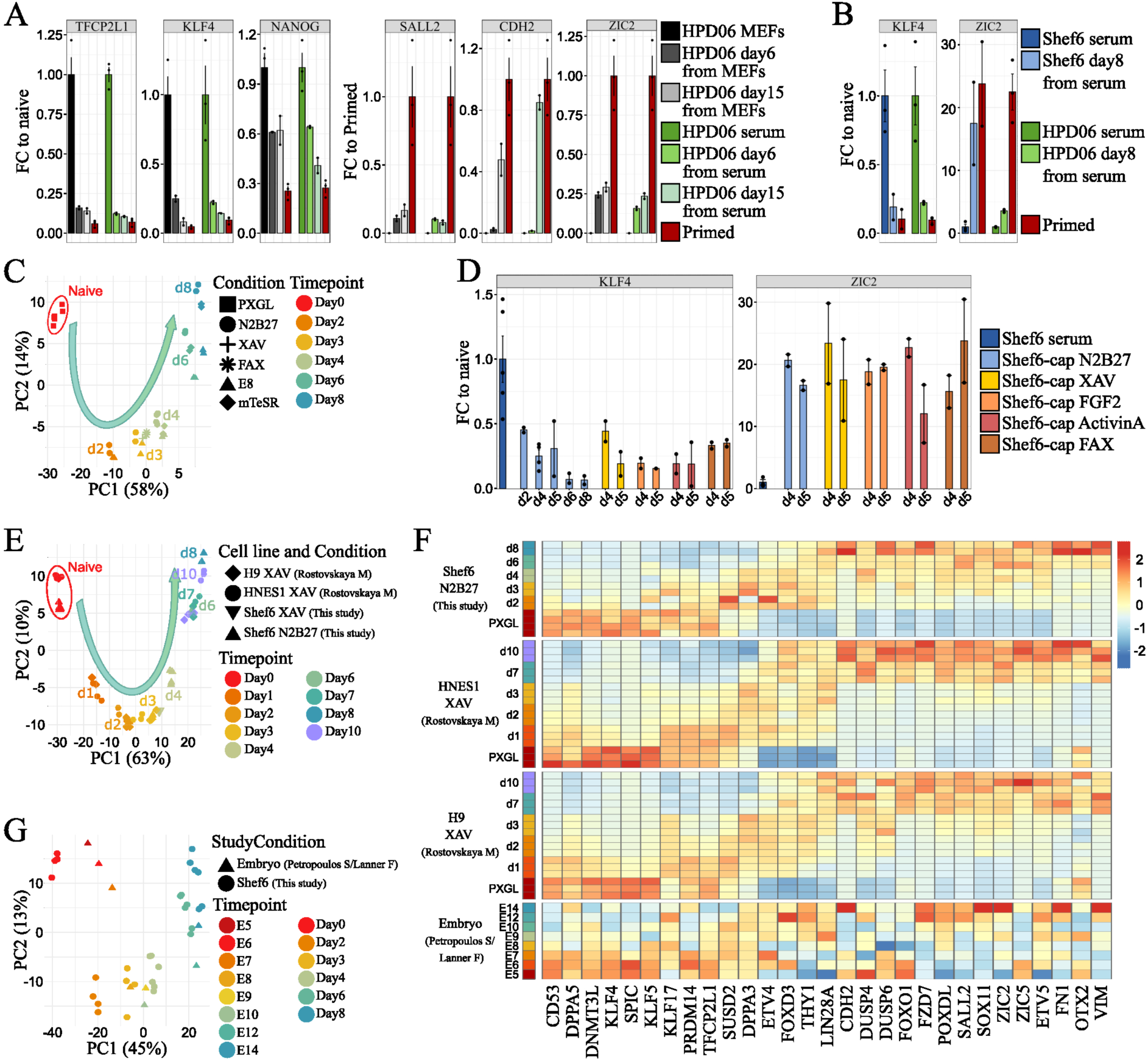
Feeder-free naive hPSCs can recapitulate embryonic differentiation trajectories. **A** Gene expression analysis by RT-qPCR of naive (*TFCP2L1* and *KLF4*), general (*NANOG*) and primed (*SALL2*, *CDH2* and *ZIC2*) pluripotency markers in HDP06 hiPSCs capacitated from naive lines stably cultured on MEFs or serum coating. Bars indicate the mean ± SEM of biological replicates shown as dots. Biological replicates include n = 3 from three independent experiments for naive hPSCs in PXGL and primed and n = 2 from two independent experiments for capacitated cells at days 6 and 15. **B** Gene expression analysis by RT-qPCR of naive (*KLF4*) and primed (*ZIC2*) pluripotency markers in Shef6 hESCs and HDP06 hiPSCs capacitated from naive lines stably cultured on serum coating. Bars indicate the mean ± SEM of biological replicates shown as dots. Biological replicates include n = 3 from three independent experiments for naive hPSCs, n = 2 from two independent experiments for capacitated cells at day 8, and n = 2 or 3 from two or three independent experiments for primed hPSCs. **C** Principal component analysis of all genes identified via RNA-seq of an eight-day differentiation time course starting from Shef6 naive hESCs stably cultured on serum coating in PXGL. Differentiation was induced by culture in basal medium (N2B27) with or without supplementation of TNKS1/2 inhibition (XAV) or FGF2-ActivinA-XAV (FAX) or commercial media containing FGF2 and Activin A signalling modulators (E8 or mTeSR). Biological replicates include n = 6 from two independent experiments for d0, from two independent experiments n = 4 for N2B27 at day 4, and n = 2 for all other combinations of conditions and time points. **D** Gene expression analysis by RT-qPCR of naive (*KLF4*) and primed (*ZIC2*) pluripotency markers of Shef6 naive hESCs stably cultured on serum coating and capacitated in basal medium (N2B27) supplemented with XAV, FGF2, Activin A or a combination of all three (FAX). Bars indicate the mean ± SEM of relative expression levels (2^−ΔΔCT^) calculated for biological replicates shown as dots. Biological replicates include n = 5 from two independent experiments for naive cells on serum, n = 4 from two independent experiments for N2B27 at day 4, and n = 2 for all other combinations of conditions and time points. **E** Principal component analysis of the abovementioned feeder-free eight-day differentiation time course in N2B27 with or without supplementation of XAV integrated with already published RNA-seq data of H9 and HNES1 hESCs cultured on MEFs and capacitated in N2B27 supplemented with XAV for up to 10 days^68^. Data was batch-corrected for data sets. Biological replicates include n = 6 from two independent experiments for d0, n = 4 from two independent experiments for N2B27 at day 4, n = 2 for all other combinations of conditions and time points, and n = 3 for each combination of cell line, condition, and timepoint in the published dataset. **F** Heatmaps showing the expression of general pluripotency, naive, and primed markers. *In vitro* data includes serum-adapted Shef6 naive hESC lines capacitated in N2B27 and published data for H9 and HNES1 hESCs cultured on MEFs and capacitated in N2B27 supplemented with XAV^68^. Biological replicates include n = 6 from two independent experiments for PXGL, n = 4 from two independent experiments for N2B27 at day 4, n = 2 for all other combinations of conditions and time points, and n = 3 for each combination of cell line, condition, and timepoint in the published dataset. The human embryo reference dataset was a subset for embryonic cells spanning E5-E14^75^. Expression levels for bulk data are row-wise z-transformed DESeq2-normalized counts. For the embryo data expression values were normalised, averaged and then log-transformed. **G** Principal component analysis of the abovementioned feeder-free eight-day differentiation time course (N2B27, E8 and mTeSR) integrated with pseudo bulk data derived from the human embryo reference dataset (subset for embryonic cells spanning E5-E14). Data was batch-corrected for data sets and then filtered by the 9376 DEGs (|log2FC| > 1, padj < 0.05) when comparing the naive state with any of the differentiation conditions and time points. Biological replicates include n = 4 from two independent experiments for d0 and n = 2 for all other combinations of conditions and time points.

To further characterise gene expression behaviour during capacitation, we explored transcriptome changes throughout an eight-day differentiation time course in various conditions: unsupplemented N2B27, N2B27 supplemented with TNKS1/2 inhibition (XAV)^68^ or FGF2-ActivinA-XAV (FAX), or E8 or mTeSR, which contain FGF and Activin signalling ligands. All differentiation was induced from naive Shef6 hESCs cultured on serum. We firstly noted that differentiation time points, rather than medium conditions, had the largest impact on clustering (Figure 4C) and that the Spearman correlation between the same time points of different conditions was high (Supplementary Figure 3C). Consistently, expression kinetics of *KLF4* and *ZIC2* were equivalent between conditions (Figure 4D). Comparison with published data for MEF-cultured H9 and HNES1 naive hESC lines differentiating with XAV^68^, showed that Shef6 naive hESCs cultured on serum coating followed the same trajectory as H9 and HNES1 hESCs cultured on MEFs before capacitation (Figure 4E), and have similar expression patterns for known pluripotency genes (Figure 4F). We also noted that the presence of XAV had no discernable effect regarding gross gene expression or proliferation compared to capacitation in N2B27 alone. We surmise that the gene expression programme accompanying capacitation is not affected by the substrate used for culture.

We next benchmarked our *in vitro* capacitation system starting from serum-coated naive hPSCs by comparing our RNA-seq time course data to data of human *ex vivo* cultured pre- and peri-implantation embryos^69^. Principal component analysis showed a striking alignment between our *in vitro* capacitation system and embryo data (Figure 4G). This is reflected in a clear correlation between differentiation-induced gene expression changes *in vitro* and *in vivo* at both the global transcriptome (Supplementary Figure 3D), and the individual pluripotency marker level (Figure 4F and Supplementary Figure 3E-F).

Taken together, these findings demonstrate that serum-adapted hPSCs progress through the developmental states of pluripotency at least as well as feeder-dependent hPSCs and that this model can generate results compatible with *in vivo* embryonic development.

### Naive hPSCs on serum coating efficiently differentiate to the three primary germ layers and TSCs

After confirming that naive hPSCs cultured on serum can exit pluripotency under different conditions with a progression similar to that of the same cell line cultured on MEFs (Figure 4 and Supplementary Figure 3), we wondered if naive hPSCs cultured in feeder-free conditions still retain the capacity to efficiently differentiate into embryoid bodies (EBs) composed of the three germ layers (Figure 1F). Capacitated hPSCs from naive cells previously cultured on MEFs or serum coating were aggregated in 3D and let spontaneously differentiate for 14 days (Figure 5A). EBs derived from both conditions showed an upregulation of germ layer markers compared to capacitated cells (Figure 5B). We conclude that naive hPSCs cultured on serum efficiently capacitate and differentiate to the three germ layers with similar or even better efficiency compared to naive hPSCs previously cultured on MEFs.

**FIGURE 5.**
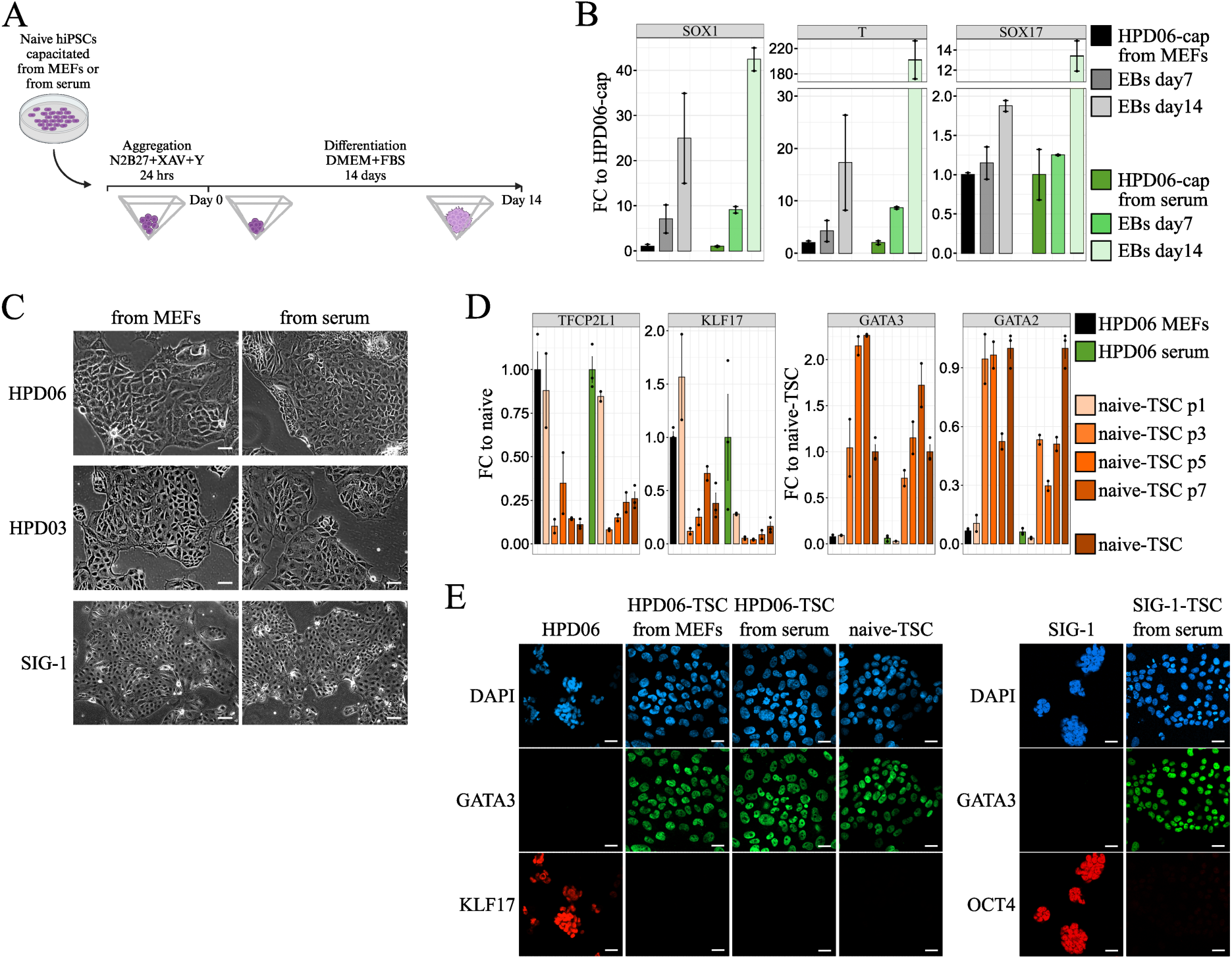
Feeder-free naive hPSCs efficiently differentiate to embryonic and extraembryonic lineages. **A** Schematic representation of the experimental setting for the EBs differentiation of HPD06 naive hiPSCs capacitated from MEFs or serum coating. **B** Gene expression analysis by RT-qPCR of germ layers markers (*SOX1*, *T*, and *SOX17*) in EBs derived from capacitated HPD06 naive hiPSCs capacitated from MEFs or serum coating. Bars indicate the mean ± SEM of n = 2 independent experiments biological replicates shown as dots. **C** Morphologies of TSCs derived from HPD06, HPD03 and SIG-1 naive hiPSCs stably cultured on MEFs or serum coating. Scale bars: 50 μm. Representative images of two biological replicates. **D** Gene expression analysis by RT-qPCR of naive pluripotency (*TFCP2L1* and *KLF17*) and TSCs (*GATA3* and *GATA2*) markers in TSCs derived from HPD06 naive hiPSCs stably cultured on MEFs or serum coating. Bars indicate the mean ± SEM of technical replicates shown as dots. Biological replicates include n = 3 independent experiments for naive hPSCs and naive-TSCs and n = 2 independent experiments for the differentiation time points. **E** Immunostaining for TSCs (GATA3) and naive (KLF17) or general pluripotency (OCT4) markers of trophectoderm derived from HPD06 and SIG-1 naive hiPSCs stably cultured on MEFs or serum coating. Scale bars: 50 μM. Representative images of two independent experiments are shown.

Naive hPSCs cultured on MEFs can also promptly differentiate toward TSCs when exposed to TSC medium^34,35,70^. Therefore, we assessed whether naive hPSCs cultured on serum coating still retain this capacity. We obtained TSCs in a few passages from naive hiPSCs stably cultured in both conditions, which showed indistinguishable morphology (Figure 4C) and similar downregulation of naive genes *TFCP2L1* and *KLF17* and upregulation of TSC genes *GATA3* and *GATA2* (Figure 5D and Supplementary Figure 4A). Consistently, the GATA3 signal was detected by immunofluorescence at a comparable level to stable naive-derived TSCs (passage 15), while KLF17 and OCT4 were undetectable (Figure 5E and Supplementary Figure 4B). Several genes (e.g. *TFAP2A*, *KRT7,* and *GATA3* and *GATA2* to a lesser extent) are shared between TSCs and amnion, as early amniogenesis has been reported to occur via a trophectoderm-like route^71^. Therefore, we wondered if TSCs derived from naive hiPSCs cultured on serum retain the same low expression of amnion markers of naive hiPSCs cultured on MEFs. We previously reported that BAP medium-treated primed hPSCs showed increased expression of *GATA3* together with amnion markers *IGFBP3* and *BAMBI*^72^. Therefore, we exposed primed Keratinocytes induced Pluripotent Stem Cells (KiPS) to BAP medium and observed a cell morphology consistent with previous reports^36,72,73^ and completely distinct from our naive hiPSCs-derived TSCs (Supplementary Figure 4C, left and Figure 4A). We analysed the expression of the amnion markers^35,70^ and detected lower expression of *IGFBP3* and undetectable expression of *BAMBI* in TSCs derived from naive hiPSCs cultured both on MEFs and serum coating compared to BAP-treated cells (Supplementary Figure 4C). We concluded that naive hiPSCs cultured in both conditions readily differentiate to *bona fide* TSCs with similar efficiency.

### Naive hPSCs on serum coating efficiently self-organize into blastoids

Previous studies have reported that naive hPSCs when aggregated are able to self-organize into blastoids. Blastoids are integrated stem cell embryo models reminiscent of the human blastocyst^40–42^. Indeed, such structures robustly specify an outer layer of TE-like cells encompassing the pluripotent epiblast (EPI)-like cells and rare PrE-like cells. To test if naive hPSCs cultured on serum retain this unique characteristic, we optimised the Kagawa *et al*^41,74^ blastoid induction protocol for naive H9 hESCs (Table 1). When passaged at high densities, naive H9 hESCs cultured on serum coating were able to aggregate in AggreWells and form blastoid structures without the need for the time-consuming plastic attachment step typically used to deplete MEFs and differentiated cells from the cultures (Figure 6A, top). Blastoids derived from naive H9 hESCs cultured on serum coating showed comparable cavitation efficiency, diameter, and rate of single cavities to blastoids derived from naive hESCs cultured on feeders (Figure 6A, top and Supplementary Figure 5A). Comparable efficiency of blastoid formation was observed also using the male naive hiPSC line KOLF2.1J cultured on MEFs or serum coating (Figure 6A, bottom). Immunofluorescence analysis validated that these structures contain TE-like (GATA2/3 or CDX2), EPI-like (NANOG or OCT4), and rare PrE-like cells (GATA4 and SOX17) (Figure 6B). Comparable results were obtained with naive HNES1 hESCs and the female naive hiPSC line SCTi003-A (Supplementary Figure 5B-C). This analysis confirmed that naive hESCs and hiPSCs cultured on serum-coated dishes are able to robustly self-organize into blastoids.

**FIGURE 6.**
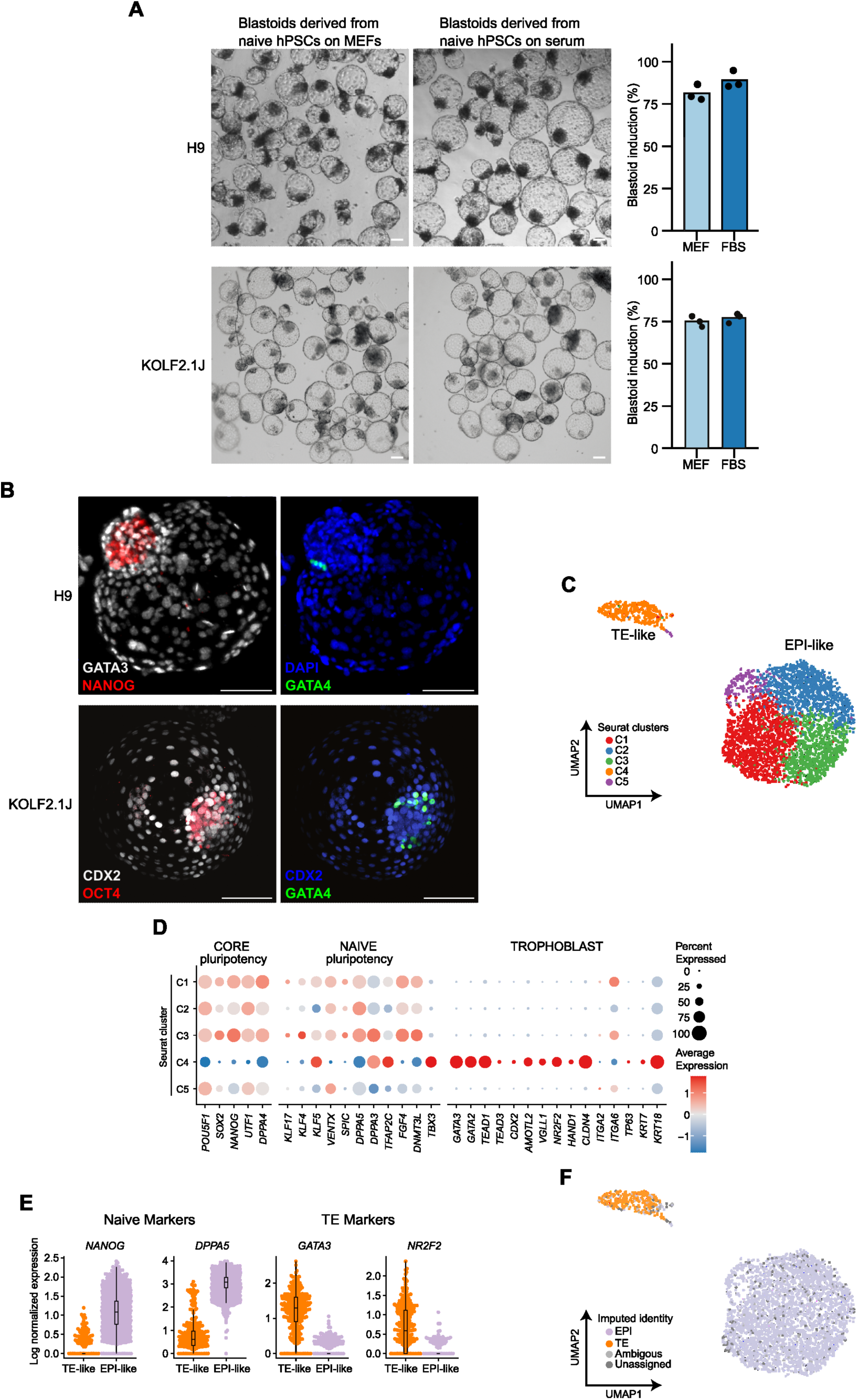
Feeder-free naive hPSCs efficiently form blastoids. **A** Left: Representative brightfield images of blastoids induced from naive H9 hESC and naive KOLF2.1J hiPSCs cultured on MEFs or serum coating. Scale bar: 100 μm. Right: quantification of induction efficiency from naive H9 hESC and naive KOLF2.1J hiPSCs cultured on MEFs or serum-coated dishes. Data is based on brightfield images of d5 blastoids. Data is presented as the mean and data points of n = 3 biological replicates for naive H9 hESC and n = 3 technical replicates for naive KOLF2.1J hiPSCs. **B** Immunostaining of d5 blastoids derived from naive H9 hESC and naive KOLF2.1J hiPSCs cultured on serum coating. Blastoids were stained for TE (GATA3 or CDX2), PrE (GATA4) and EPI (NANOG and OCT4) markers. Shown is the maximum projection. Scale bars: 100 μm. **C** UMAP plot from scRNA-seq analysis of blastoids from naive H9 hESCs cultured on serum coating. Cells are coloured according to the sample Seurat cluster (n = 3260). Highlighted are TE-like (cluster 4) and EPI-like clusters. **D** Expression of selected lineage-specific marker genes across Seurat clusters from (C). The size of the dots represents the proportion of cells in the indicated group expressing the given gene and colour encodes the scaled average expression. **E** Combined beeswarm and box plots of gene expression of naive pluripotency markers (*NANOG* and *DPPA5*) and TE markers (*GATA3* and *NR2F2*) from (C). Cells were separated based on their EPI or TE signature. Single cells are plotted as dots; boxes span the interquartile range (IQR), the centre line indicates the median and whiskers extend to values within 1.5 times the IQR. **F** UMAP plots as in (C) but with cells based on the imputed annotation of *in vitro* samples from a reference in vivo dataset^69^. Unassigned and ambiguous labels refer to cells with either none or with more than two imputed annotations respectively.

To validate these cellular identities, we performed 10x scRNAseq on day 5 blastoids derived from naive H9 hESCs cultured on serum-coated dishes. This analysis showed two populations of cells on the UMAP (Figure 6C). Based on the expression of key marker genes, the two populations were annotated as TE-like and EPI-like cells (Figure 6D-E). Notably, TE-like cells expressed a wide range of trophoblast markers (e.g., *GATA3, NR2F2, GATA2*) but not those of the amnion (e.g., *ISL1, TFAP2B*) or other off-target lineages (Figure 6D-E, Supplementary Figure 5D). Similarly, EPI-like cells expressed both core (e.g., *POU5F1, NANOG*) and naive pluripotency markers (e.g., *DPPA5, DNMT3L*) but not those associated with primed pluripotency (e.g., *OTX2, ZIC2*) (Figure 6D-E, Supplementary Figure 5D). Previous reports have shown that PrE-like cells are very rarely specified during blastoid induction^41^. Consistent with this finding, we only observed sporadic individual cells expressing the PrE marker *GATA4*, and no Seurat cluster contained a significant number of PrE-like cells (Supplementary Figure 5D-E). To further validate these annotations, we integrated our data with embryo reference datasets (Supplementary Figure 5F)^20,40,69,75–78^. Notably, our single-cell dissociation protocol disproportionately fragilises the TE-like cells; therefore, scRNA-seq was only used to verify cellular identities rather than to quantify the proportion of individual lineages. Nevertheless, our analysis confirmed that cluster 4 is predominantly composed of TE-like cells, while the remaining clusters retain EPI identity (Figure 6F, Supplementary Figure 5F-G). This analysis confirmed that the blastoids contain *bona fide* TE-like and EPI-like cells. Overall, the formation of blastoids was achieved using 4 lines in 3 different laboratories, demonstrating the robustness of the protocols and their potential for wide adoption in the community.

Overall, feeder-free naive hESCs, when aggregated, robustly self-assemble into blastoids containing cells with appropriate lineage identities.

## Discussion

Expansion of human naive PSCs on a layer of inactivated MEFs has several limitations. Production of MEFs requires the sacrifice of mice, which contrasts with the 3Rs principles for the ethical use of animals in research. MEF production is also time-consuming and expensive, limiting the range of application of human naive PSCs. Indeed, both mESCs and conventional hPSCs became widespread models only once feeder-free conditions were developed^3,13–15^.

The use of MEFs brings an additional hurdle, which is the co-culture of non-human cells with hPSCs. The former might appear as contaminants in global analyses of the latter, as suggested by our analyses of several RNA-sequencing datasets revealing widespread expression of fibroblast markers (Supplementary Figure 2C). Furthermore, the two cell populations might affect each other in a non-controllable manner.

Conventional hPSCs can be expanded without feeders on plates coated with recombinant ECM proteins, such as fibronectin and vitronectin^14,15^. Alternatively, solubilised basement membrane matrixes, commercialised as Matrigel or Geltrex, can be used^13^. However, these substrates have been tested for human naive PSC expansion with only limited success. In some cases (i.e. laminin) only short-term expansion could be achieved, while in others (i.e. Matrigel) stable expansion could be achieved only with additional inhibitors in the media^23,24,53^.

Importantly, these substrates are expensive and range from 0.5 to 2 Euros/cm^2^. In contrast, the cost of using serum coating is reduced by two orders of magnitude to around 0.5 cents/cm^2^, while allowing for long-term expansion without requiring any additional changes in media formulation or culture regime. For all these reasons, serum-coating represents a significant improvement over current methodologies, allowing for the widespread use of naive hPSCs for several applications, such as large-scale screenings, epigenetic and metabolic profiling, and the generation of embryo models such as blastoids. Expansion of naive hPSCs on serum resulted in a streamlined blastoid protocol, which worked in multiple laboratories with multiple cell lines, with an efficiency as high as feeder-based protocols.

A recent study showed that human naive PSC colonies are surrounded by laminins and collagens^52^, which are, at least in part, produced by naive hPSCs themselves. However, this autocrine ECM protein production is not sufficient for PSC expansion, as either MEFs or exogenous substrates are needed. Our proteomic analysis of the FBS coating allowing for human naive PSC expansion revealed the presence of several Collagens, Laminins, together with Vitronectin and Fibronectin. We, therefore, speculate that this combination of ECM proteins is crucial for naive hPSC expansion, but future functional studies will be needed to tease out the contribution of each individual ECM protein.

Implementing our approach requires batch testing of serum, a standard procedure for over 40 years, also required for the culture of mESCs with serum-based media^79^. Indeed, in our experience, FBS batches allowing robust expansion of mouse ESCs all worked well for providing growth substrates for naive hESCs.

We also observed that harsh treatments, such as growth at clonal density after fluorescence-activated cell sorting in the presence of antibiotics are better tolerated on feeders. Thus, some procedures, such as genomic engineering, might be performed preferably on MEFs. Genomic engineering on mESCs via homologous recombination has been performed on feeders for similar reasons^80–84^. We note that shuttling serum-adapted hESCs between serum coating and MEFs works without the need for re-adaptation to feeder-free conditions.

Finally, this manuscript represents a joint effort of several laboratories routinely working with naive hPSCs. Once a methodology was identified by one lab, reagents and protocols were shared with other labs, resulting in extensive testing of multiple lines for different applications before publication. This constructive and cooperative approach allowed us to quickly build trust in our methodology, thanks to the testing performed in parallel by 5 labs.

## Materials and Methods

### Culture of hPSCs

Naive hiPSCs HPD06 and HPD03 were previously generated by direct reprogramming from somatic cells as described in^30^ were cultured on mitotically inactivated mouse embryonic fibroblasts (MEFs; DR4 ATCC) or serum coating in PXGL medium ^32^ at 37°C, 5% CO_2_, 5% O_2_. The PXGL medium was prepared as follows: N2B27 (DMEM/F12 [Gibco 11320074], and Neurobasal in 1:1 ratio [Gibco 21103049], with 1:200 N2 Supplement [Gibco 17502048], and 1:100 B27 Supplement [Gibco 17504044], 2 mM L-glutamine [Gibco 25030024], 0.1 mM 2-mercaptoethanol [Merck Sigma Aldrich M3148]) supplemented with 1 µM PD0325901 (Axon Medchem 1408), 2 μM XAV939 (Axon Medchem 1527), 2 μM Gö6983 (Axon Medchem 2466) and 10 ng/ml human LIF (Qkine Qk036). Serum coating medium was composed of cold 10% Fetal Bovine Serum [Gibco A5256701 or Sigma-Aldrich F7524 or Biowest S1600] in DMEM high glucose [Gibco 41965039]. After adding 2ml of serum coating solution to each well of a 6-well plate, plates were incubated overnight at 37°C, rinsed once with PBS without MgCl2/CaCl2 (Merck Sigma Aldrich D8537) before plating the cells. Cells were passaged as single cells every 3-4 days at a split ratio of 1:3 or 1:4 following dissociation with TrypLE (Gibco 12563029) for 3 minutes at room temperature. 10 μM ROCK inhibitor (ROCKi, Y27632, Axon Medchem 1683) was added to the naive medium for 24 hours after passaging. For the feeder-free conversion, naive hiPSCs HPD06 and HPD03 stably cultured on MEFs were collected with ReLeSR (STEMCELL Technologies 100-0483). Cells were first incubated for 2 minutes at room temperature, followed by another 5 minutes at 37°C after the complete aspiration of the reagent. Cells were collected by carefully washing the well with fresh PXGL medium adding 10 µM ROCKi and were plated on feeder-free plates coated with serum at a 1:1 to 1:2 ratio.

Naive hESCs H9^1^ from Leeb’s group and Shef6^85^ were cultured on mitotically inactivated DR4 MEFs or serum coating (DMEM high glucose [Sigma-Aldrich D5671] and 10% FBS (Sigma-Aldrich F7524 or Biowest S1600) in PXGL at 37°C, 5% CO_2_, 5% O_2_. Cells were passaged every 3-4 days at a split ratio up to 1:12 using Accutase (Merck Sigma Aldrich A6964) for 5 minutes at 37°C. 10 μM ROCK inhibitor was added to the naive medium for 24 hours after passaging. Naive H9 hESCs from Zylicz’s group were instead passaged at a split ratio up to 1:6 using TrypLE (Gibco 12563029) for 3 minutes at 37°C.

Naive KOLF2.1J^86^ and SCTi003-A (STEMCELL Technologies 200-0511) hiPSCs from Rivron’s group were cultured on 0.1% gelatin-coated plates including a feeder layer of irradiated MEFs or serum coating (DMEM high glucose and 10% FBS (Gibco A5256701)) in PXGL at 37°C, 5% CO2, 5% O2. Cells were passaged every 3-4 days in a ratio of 1:6 following dissociation with Accutase (Merck Sigma Aldrich A6964) for 5 minutes at 37°C. 10 μM ROCK inhibitor was added for 24 hours after passaging. Naive H9 hESCs from Pasque’s, SIG-1 hiPSCs (Sigma-Aldrich iPSC EPITHELIAL-1-IPSC0028), and HNES1 hESCs^26^ were cultured on MEFs or serum coating (10% Fetal Bovine Serum [Gibco 10270] in DMEM high glucose [Gibco 41966029]) in PXGL at hypoxia conditions in 5% O2 and 5% CO2 incubator under humidified conditions at 37°C. Cells were passaged every 3 days at a split ratio up to 1:10 by single-cell dissociation with Accutase (Sigma-Aldrich, A6964-100ML) for 5 minutes at 37°C. 10 μM ROCK inhibitor was added to the naive medium for 24 hours after passaging.

Human primed KiPS^24^ were cultured on pre-coated plates with 0.5% growth factor-reduced Matrigel in E8 medium, made in-house according to ^15^, at 37°C, 5% CO2, 5% O2. Cells were passaged every 3– 4 days at a split ratio of 1:8 following dissociation with 0.5 mM EDTA (Invitrogen AM99260G) in PBS without MgCl_2_/CaCl_2_.

Primed hESCs H9 from Leeb’s group (WiCell#WA09) and re-primed Shef6 were maintained feeder-free on plates pre-coated with 1 % (v/v) Geltrex: (Gibco A1413302) in E8 flex (Gibco A2858501) supplemented with 1 % (v/v) Pen/Strep at 37°C, 5% CO2, 5% O2. Following dissociation with Versene (Gibco 15040-033) at room temperature for 5 minutes, cells were passaged every 3-4 days at a split ratio of 1:6.

Primed hESCs H9 from Pasque’s group were grown in pre-coated geltrex tissue culture at normoxia conditions and under humidified conditions at 37°C in complete E8Flex medium (Thermo Fisher, A2858501). Cells were dissociated into clumps every 5–6 days by incubating 5 min at room temperature in Accutase and passaged at a split ratio of 1:12. 10 μM ROCK inhibitor was added to the naive medium for 24 hours after passaging.

All cell lines were routinely checked for mycoplasma and tested negative (Euroclone EMK090020 or Mycoplasma check Service from Eurofins).

### Capacitation of naive hPSCs

Naive hiPSCs HPD06 stably cultured on MEFs or serum coating were capacitated following Rostovskaya *et al*^68^ protocol with minor modifications. Briefly, cells were dissociated with TrypLE, and 5×10^4^ cells/cm^2^ were seeded on a 6-well plate pre-coated with 0.5% growth factor-reduced Matrigel in E8 medium with 10 µM ROCKi (day 0). From day 1 to day 6 cells were cultivated in N2B27 with 2 μM XAV939. From day 6 onward the medium was changed to XAF (N2B27 supplemented with 2 μM XAV939, 20 ng/ml Activin A [Qkine Qk001] and 10 ng/ml FGF2 [Qkine Qk002]).

Naive hESCs Shef6 and H9 stably cultured on MEFs or serum coating were capacitated following a modified version of the Rostovskaya *et al*^68^ protocol. Briefly, cells were dissociated with Accutase and 1×10^4^ cells/cm^2^ or 1.5×10^4^ cells/cm^2^ were seeded on a 6-well plate pre-coated with Geltrex (Thermo Fisher Scientific A1413302) directly into N2B27 medium supplemented with 10 µM ROCKi (day 0). From day 1, cells were cultivated in N2B27 without ROCKi. For the additional conditions, the N2B27 was supplemented with 2 µM XAV939, 12 ng/mL FGF2 (in-house or PeproTech 100-18B) and/or 10 ng/mL Activin A (PeproTech 120-14E) or replaced with E8 or mTeSR.

### Conversion of human naive PSCs into trophectoderm/TSCs

Naive hiPSCs HPD06 and HPD03 stably cultured on MEFs or serum coating were pre-treated on MEF with TS medium for 24h. Cells were dissociated with TrypLE, and 5×10^4^ cells/cm^2^ were seeded on a 6-well plate pre-coated with 5 μg/mL Collagen IV and further cultured in TS medium (DMEM/F12 supplemented with 0.1 mM 2-mercaptoethanol, 0.2% FBS, 0.3%, Bovine serum albumin (BSA, Gibco 15260-037), 1% ITS-X (Gibco 51500), 1.5 μg/ml L-ascorbic acid (Merck Sigma Aldrich A4544), 50 ng/ml EGF (Qkine Qk011), 2 μM CHIR99021 (Axon Medchem 1386), 0.5 μM A83-01 (Axon Medchem 1421), 1 μM SB431542 (Axon Medchem 1661), 0.8 mM VPA [HDACi, Merck Sigma Aldrich P4543], and 5 μM ROCKi). Cells were cultured in 5% CO_2_ and 5% O_2_ changing medium every 2 days and were passaged upon 80% confluency at a 1:6 to 1:8 ratio.

Naive SIG-1 hiPSCs to trophoblast differentiation was done as previously published^39^, using the following previously described protocols for hTSCs^34–36,70^. Cells were cultured on MEFs or serum coating until subconfluency and dissociated in single-cell using TryplE for 5 min at 37°C in PXGL supplemented with 10 mM Y-27632 (Tocris, 1254). At day 2 of culture, cells were washed with PBS (Gibco, 10010-015), and the media was switched from PXGL to ASECRiAV medium^87^ consisting of DMEM/F12 (Gibco, 31330038) supplemented with 0.3% BSA (Sigma A3059), 0.2% FBS (Gibco 10270-106), 1% Penicillin-streptomycin (Gibco 15140-122), 1% insulin-transferrin-selenium-ethanolamine-X 100 supplement (Gibco 51500056), 1.5 mg/mL L-ascorbic acid (Sigma A8960), 0.5 mM A83-01 (Peprotech 9094360), 1 mM SB431542 (Axon Medchem 1661), 50 ng/ml hEGF (Miltenyi Biotec 130-097-750), 2uM CHIR99021 (Axon Medchem 1386), 0.8 mM Valproic acid (Sigma V0033000), 0.1 mM b-mercapto-EtOH (Gibco 31350-010) and 5 mM Y-27632 (Tocris 1254). The medium was changed every day and supplemented with 5 mM Y-27632. From passage 1 onwards hTSCs were cultured and maintained on 1.5 mg/mL iMatrix-silk-E8-Laminin overnight coated tissue culture treated plates in hypoxia conditions 5% O2 and 5% CO2 at 37°C and passaged every 5 days at 1:3 splitting ratio. hTSCs used in all experiments were cultured and maintained in ASECRiAV until day 26 of conversion.

Naive SIG-1 hiPSCs differentiation to trophectoderm was performed as previously described^88^. Briefly, cells were dissociated in single cells using Accutase at 37°C for 6 min, followed by gentle mechanical dissociation. Single cells were counted and viability was assessed via trypan blue staining using ThermoFisher Countess II and 3×10^4^ live cells/cm^2^ were plated onto iMatrix Silk laminin 511-coated plate (Amsbio AMS.892 012) in nTE1 media (N2B27 supplemented with 2 µM A83-01 [Merck Sigma Aldrich SML0788], 2 µM PD0325901 [Axon Medchem 1408], 10 ng/ml BMP4 [R&D Systems 314-BP] and 10 µM Y-27632 [Tocris 1254]). After 24h, cells were washed with PBS and media switched to nTE2 for 48h and refreshed daily (N2B27 supplemented with 2 µM A83-01, 2 µM PD0325901 and 1 µg/ml JAKi1 [Merck Sigma Aldrich 420099]). Cells were fixed on day 3.

### Culture of BAP-treated primed pluripotent stem cells

MEFs were fed with DMEM/Ham’s F-12 medium containing 0.1 mM 2-mercaptoethanol, 1% Insulin-Transferrin-Selenium-Ethanolamine (ITS-X; Gibco 51500056), 1% non-essential amino acids (NEAA; Gibco 11140050), 2 mM L-glutamine, and 20% KnockOut™ Serum Replacement (KSR; Gibco 10828028) for 24 h. The supernatant was collected and used as MEF-conditioned medium. KiPS were dissociated into single cells with TrypLE. The cells were cultured on pre-coated plates with 0.5% growth factor-reduced Matrigel at a density of 2 × 10^4^ cells/cm^2^ with MEF-conditioned medium supplemented with 10 ng/ml BMP4 (Peprotech 120-05ET), 1 μM A83-01, 0.1 μM PD173074 (Axon Medchem 1673), and 10 μM ROCKi, as described previously^36^. The medium was changed daily.

### Spontaneous differentiation with embryoid bodies

Capacitated hiPSCs HPD06 from stable cultures on MEFs or serum coating were single-cell dissociated with TrypLE and seeded at 1.5×10^4^ cells in v-bottom 96-well plates (Greiner, M9686) pre-treated with anti-adherence AggreWell™ Rinsing Solution (STEMCELL Technologies 7010) in E8 medium with 10 µM ROCKi (day 0). From day 1, aggregates were cultivated in spontaneous differentiation medium (DMEM/F12 with 20% FBS, 2 mM L-glutamine, 1% NEAA and 0.1 mM 2-mercaptoethanol). Half of the medium was changed every 2 days.

### Blastoid formation

Blastoid experiments from naive H9 hESCs were performed according to Kagawa *et al*^41^, with minor modifications. Briefly, H9 naive hESCs were collected by incubation for 3 min with Accutase (Merck Sigma Aldrich A6964). Single cells were plated at a density of 98-102 cells per micro-well of a 24-well AggreWell^TM^400 plate in N2B27 supplemented with 10 μM Y27632 (Axon Medchem 1683) to aggregate for 3h. Subsequently, half medium was changed with 2x PALLY consisting of N2B27 supplemented with 1 μM PD0325901 (Axon Medchem 1408), 1 μM A83-01 (Axon Medchem 1421), 10 ng/mL hLIF (Qkine Qk036), 1 μM oleoyl-L-α-lysophosphatidic acid sodium salt (LPA, Merck Sigma Aldrich L7260) and Y27632. 24 hours later, half-medium was changed with 1x PALLY. After 48h of PALLY culture, blastoids were maintained for 56 hours in LY medium consisting of N2B27 supplemented with LPA and Y27632. Blastoids from H9 naive hESCs were processed for single-cell RNA sequencing at day 5.

Blastoid experiments from naive KOLF2.1J and SCTi003-A hiPSCs were performed according to Kagawa et al.^41^, with minor modifications. Naive KOLF2.1J and SCTi003-A hiPSCs were collected by incubation with Accutase for 5 min. In the presence of MEFs, cells were incubated for 70 minutes on 0.1% gelatin-coated plates after which the non-attached hiPSCs were collected. Single cells were plated at a density of 20.000 cells per well of a 96-well plate comprising microwells of 200 µm diameter, made as previously described^89^. Cells were cultured in N2B27 supplemented with 0.3% BSA (Sigma Aldrich A7979), 1µM PD0325901 (MedChem Express HY-10254), 1µM A83-01 (MedChem Express HY-10432), 10ng/ml hLIF (in house), 1 µM LPA (Peprotech 2256236) and 1:1000 CEPT cocktail (Thermo Fisher A56799). After 24 hours the medium was changed to N2B27 supplemented with 1 µM PD0325901, 1 µM A83-01, 10 ng/ml LIF and 1 µM LPA for 48 hours with daily medium changes. Blastoids were maintained for an additional 48 hours in N2B27 medium supplemented with 1 µM LPA. Blastoid experiments from naive HNES1 hESCs were performed as previously described^41^ using AggreWells™400 (STEMCELL Technologies 34415). Briefly, naive HNES1 hESCs were treated with Accutase at 37°C for 6 min, followed by gentle mechanical dissociation. Cell pellet was resuspended in Aggregation medium composed of N2B27 with 0.3% BSA (Merck Sigma Aldrich A7979) and 10 µM Y-27632. Single cells were counted and viability assessed via trypan blue staining using ThermoFisher Countess II and 1.0/1.2×10^5^ live cells/well (∼80/100 cells/μwell) were seeded into AggreWells™400 and grown in a hypoxic chamber. AggreWells™400 were treated with an anti-adherent solution (STEMCELL Technologies 07010) and washed 3 times (2x N2B27 and 1x aggregation medium) before cell plating (day −1). The cells were allowed to form aggregates inside the microwell for 24 hours. On day 0, the aggregation medium was replaced with PALLY medium: N2B27 with 0.3% BSA, 1 µM PD0325901, 1 µM A83-01 (Merck Sigma Aldrich SML0788), 0.5 µM LPA (Tocris 3854), 10 ng/ml hLIF and 10 µM Y-27632. PALLY medium was refreshed on day 1. After 48 h (day 2) PALLY medium was replaced with N2B27 containing 0.3% BSA, 0.5 µM LPA and 10 µM Y-27632 (LY medium). LY medium was refreshed on day 3, and blastoids fully formed on day 4.

### Proliferation assay and AP staining

Cell proliferation was assessed by plating 5×10^4^ cells/cm^2^ in a 6-well plate. Cells were single-cell dissociated with TrypLE, counted and re-plated every 3 days.

For AP staining, cells were fixed with a citrate–acetone–formaldehyde solution and stained using the Alkaline Phosphatase kit (Merck Sigma Aldrich 86R-1KT). Plates were scanned using a Nikon Scanner and scored with the Analyse Particles plugin of the software Fiji (from ImageJ) 1.0 (pixels 5-500, circularity 0.5-1).

### Gene expression analysis by quantitative PCR with reverse transcription

Total RNA from naive hiPSCs HPD06 and HPD03 was isolated using Total RNA Purification Kit (Norgen Biotek 37500), and complementary DNA (cDNA) was made from 500 ng using M-MLV Reverse Transcriptase (Invitrogen 28025-013) and dN6 primers. Data were acquired with the Applied Biosystems QuantStudio™ 6&7 Flex Software 1.3.

Total RNA from naive hESCs H9 and Shef6 was isolated with the ExtractMe kit (Blirt EM15) following the manufacturer’s instructions, and cDNA was made from 100 ng using the SensiFAST cDNA Synthesis Kit (Bioline BIO-65054). Real-time PCR was performed on the CFX384 Touch real-time PCR detection system (Bio-Rad).

SYBR Green Master mix (Bioline BIO-94020) was used for real-time qPCR and primers are listed in Supplementary Table 1. Three technical replicates were carried out for each biological replicate of all RT-qPCR analyses and *β-ACTIN* or *GAPDH* were used as endogenous controls to normalise expression.

### Immunofluorescence

Immunofluorescence analysis of naive hiPSCs HPD06 and HPD03 was performed on 1% Matrigel-coated glass coverslip in wells. Cells were fixed in 4% Formaldehyde (Merck Sigma Aldrich 78775) in PBS for 10 min at room temperature, washed in PBS, and permeabilised and blocked in PBS with 0.3% Triton X-100 (PBST) and 5% of horse serum (ThermoFisher 16050-122) for 1 hour at room temperature. Cells were incubated overnight at 4°C with primary antibodies in PBST with 3% horse serum. After washing with PBS, cells were incubated with secondary antibodies (Alexa, Life Technologies) for 45 min at room temperature. Nuclei were stained using Fluoroshield™ with DAPI (Merck Sigma Aldrich F6057). Images were acquired with a Zeiss LSN700 confocal microscope using ZEN 2012 software. Images were processed using the software Fiji (from ImageJ) 1.0. Fluorescence intensity was quantified using Cell Profiler software (v4.2.1). DAPI staining was used to identify individual nuclei of cells. At least 700 nuclei from five randomly selected fields from two independent experiments were analysed for each condition. Single-cell measured mean intensity values were considered for downstream data analysis.

For immunofluorescence analysis of naive H9 hESCs and Shef6 hiPSCs from Leeb’s group, cells were plated onto µ-Slide 8 Well Glass Bottom Chamber Slides (Ibidi 80827) with MEFs, or onto slides coated with serum or Geltrex. Cells were fixed in 4% Formaldehyde in PBS for 20 min at room temperature, washed in PBS, and permeabilised and blocked in PBS with 0.5 % Triton X-100 (PBST) and 5% BSA for 30 min at room temperature. Cells were incubated overnight at 4°C with primary antibodies in PBST with 5% BSA. After washing with PBS, cells were incubated with secondary antibodies (Alexa, Life Technologies) for 1h at room temperature. After washing with PBS, the nuclei were stained with 20 ng/mL DAPI in PBS (Invitrogen D1306) for 5 min in the dark at room temperature. Samples were mounted using Vectashield with DAPI (Vector Laboratories VECH-1200). Images were acquired with an Axio Observer Z1 microscope and processed using the software Fiji (from ImageJ) 1.0.

Immunofluorescence staining protocol was performed in naive H9 hESCs from Pasque’s group and SIG-1 hiPSC as previously published^39,90^. Cells were grown on 0.1% gelatinised 18 mm round coverslips on MEFs or serum coating. Cells were fixed in 4% paraformaldehyde-PBS for 10 min at room temperature in the dark and permeabilised with 0.5% Triton X-100 in PBS for 5 min and washed twice with 0.2% Tween 20 in PBS (PBST) for 5 min each before proceeding to the staining. Primary and secondary antibodies were diluted in a blocking buffer containing mainly 2% Tween-PBS with 5% normal donkey serum and 0.2% fish skin gelatin. Cells on coverslips were incubated overnight at 4°C with the specific primary antibodies in blocking solutions (1:50 dilution for NANOG, 1:300 dilution for KLF17), then washed three times with PBST for 5 min. After washing, the samples were incubated with secondary antibodies diluted in blocking buffer for 1 hour in the dark at room temperature. The samples were then washed with PBST three times for 5 min each and afterwards washed with 0.002% DAPI (Sigma-Aldrich D9542) solution in PBST. The coverslips were mounted in Prolong Gold reagent with DAPI after a final wash in PBST. Naive hPSCs immunofluorescence images were taken in a Nikon TiE A1R inverted microscope and were processed using the software Fiji (from ImageJ) 1.0.

Blastoids from naive H9 hESCs were fixed with 4% paraformaldehyde for 20 min at room temperature. Subsequently, the samples were washed for 3 times 10 minutes with PBST (PBS containing 0.1% Tween20) supplemented with BSA (0.3%). The samples were then permeabilised for 20 min using 0.2% Triton X-100 in PBS and afterwards blocked using a blocking buffer containing 0.1% Tween20, 1% BSA and 10% normal donkey serum in PBS for at least 3 hours. The samples were then incubated overnight at 4°C with primary antibodies diluted in blocking buffer. The next day, samples were washed with PBST at least three times for 10 min each. After washing, the samples were incubated with secondary antibodies diluted in blocking buffer for 3 hours in the dark at room temperature. The samples were then washed with PBST three times for 10 min each and afterwards mounted using PBS. Image acquisition was performed using a Leica Stellaris 5 fluorescence confocal microscope and were processed using the software Fiji (from ImageJ) 1.0.

Blastoids from naive KOLF2.1J and SCTi003-A hiPSCs were fixed with 4% PFA for 30 min at room temperature and rinsed three times with PBS. Samples were permeabilised and blocked using 0.3% Triton X-100 and 10% normal donkey serum in PBS for at least 60 min and incubated overnight at room temperature with primary antibodies diluted in fresh blocking/permeabilisation buffer. The following day samples were washed with PBS containing 0.1% Triton X-100 (PBST) at least three times for 10 min each. Samples were then incubated in Alexa Fluor-tagged secondary antibodies (Abcam or Thermofisher Scientific) diluted in PBST for at least 30 min in the dark at room temperature and washed three times with PBST for 10 min each. Images were captured using an Olympus IX83 confocal microscope. Each image consisted of 30 optical sections, with an average thickness of approximately 3–5µm per section. Images were processed using the software Fiji (from ImageJ) 1.0. Blastoids from naive HNES1 hESCs were fixed with 4% PFA for 30 min at room temperature and rinsed three times with PBS. About 10-20 structures per condition were then moved to Thermo Scientific™ Nunc™ MiniTrays with Nunclon™ Delta surface and permeabilised for 30 min at RT using PBS/0.3% Triton X-100 (Merck Sigma Aldrich). Cells were blocked for 4-6h at RT with blocking solution (PBS 0.3% Triton X-100 with 10% Normal Donkey Serum [Merck Sigma Aldrich S30]). Primary antibodies (listed in Supplementary Table 2) were diluted in blocking solution and incubated overnight at 4°C in a humidified chamber. Cells were washed three times with PBS 0.1% Triton X-100, and stained with secondary antibodies for 1h at RT in a humidified chamber. Cells were washed three times with PBS 0.1% Triton X-100, and stained with DAPI for 15 min at RT during the second wash (0.1 µg/ml DAPI in PBS 0.1% Triton X-100). Finally, blastoids were moved into IBIDI µ-Slide 15 Well 3D Glass Bottom (81507) and imaged using a C2 or TiE A1R confocal microscope (Nikon). Images were processed using the software Fiji (from ImageJ) 1.0, Z-stacks were shown as Max Intensity projections and denoised using the “despeckle” tool equally on all images.

All used primary antibodies are listed in Supplementary Table 2.

### Flow cytometry

For flow cytometry analysis, naive and differentiating Shef6 hESCs and primed hPSCs were detached using Accutase and resuspended in DMEM supplemented with 0.5% BSA. For surface marker staining the cells were washed 1x in FACS buffer (1x PBS and 1% BSA), stained with APC-conjugated anti-human SUSD2 antibody (BioLegend 327408) diluted 1:100 in FACS buffer for 30 min on ice in the dark and washed again 1x with FACS buffer. For live/dead discrimination 5 µg/mL DAPI (Invitrogen D1306) was used. Antibody signal levels were measured using the ZE5 (Bio-Rad) and then analysed with the FlowJo software (v10.10, BD Bioscience).

Nave H9 hESCs and SIG-1 hiPSCs were first stained for viability in 1:300 Live/Death Zombie Aqua UV in PBS. Fluorophore conjugated antibody was diluted at a ratio of 1:50 in fluorescence-activated cell sorting (FACS) buffer with PE-conjugated anti-human SUSD2 (Biolegend, 327406), for 20 minutes at room temperature, then washed with FACS buffer, and fixed in 4% paraformaldehyde. Cells were passed through a 40 µm cell strainer (Corning 352340). The fluorescence intensity of 20,000 cells was recorded on a flow cytometer (Symphony A5, BD Biosciences) and analyzed using FlowJo software (v10.10, BD Bioscience). Single-stained controls were used for compensation and gating in the flow cytometer. Only lived cells were used for data analysis. The frequency of the parent population has been used to plot the SUSD2-positive cells.

### Proteomics

Samples were processed using the PreOmics iST sample preparation kit (PreOmics P.O.00027). LC-MS/MS analysis consisted of a NanoLC 1200 coupled via a nano-electrospray ionization source to the quadrupole-based Q Exactive HF benchtop mass spectrometer^91^. For the chromatographic separation, a binary buffer system consisting of solution A: 0.1% formic acid and B: 80% acetonitrile, 0.1% formic acid was used. Peptides were separated according to their hydrophobicity on an analytical column (75 μm) in-house packed with C18-AQ 1.9 μm C18 resin with a gradient of 7–32% solvent B in 45 min, 32–45% B in 5 min, 45–95% B in 3 min, 95–5% B in 5 min at a flow rate of 300 nl/min. MS data acquisition was performed in DIA (Data Independent Acquisition) mode using 32 variable windows covering a mass range of 300–1650 m/z. The resolution was set to 60’000 for MS1 and 30’000 for MS2. The AGC was 3e6 in both MS1 and MS2, with a maximum injection time of 60 ms in MS1 and 54 ms in MS2. NCE was set to 25%, 27.5%, 30%. All acquired raw files were processed using Spectronaut software (17.0). For protein assignment, spectra were correlated with the Rattus Norvegicus/Bos Taurus database (v. 2023). Searches were performed with tryptic specifications and default settings for mass tolerances for both MS and MS/MS spectra.

The other parameters were set as follows. Fixed modifications: Carbamidomethyl (C); variable modifications: Oxidation, Acetyl (N-term); digestion: Trypsin, Lys-C; min. peptide length = 7Da; max. peptide mass = 470Da; false discovery rate for proteins and peptide-spectrum = 1%. Perseus software (1.6.2.3) was used to logarithmise, group and filter the protein abundance. The obtained list of proteins was analysed using the Matrisome AnalyseR tool (https://sites.google.com/uic.edu/matrisome/tools/matrisome-analyzer^62^) from The Matrisome Project.

### Exome sequencing

NEGEDIA S.r.l. performed the Exome Sequencing service. Genomic DNA was extracted with a paramagnetic beads technology using the Negedia DNA extraction service, quantified using the Qubit 4.0 fluorimetric Assay (Thermo Fisher Scientific) and sample integrity, based on the DIN (DNA integrity number), was assessed using a Genomic DNA ScreenTape assay on TapeStation 4200 (Agilent Technologies). Libraries were prepared from 100 ng of total DNA using the NEGEDIA Exome sequencing service which included library preparation, target enrichment using Agilent V8 probe set, quality assessment with FastQC v0.11.9 (https://www.bioinformatics.babraham.ac.uk/projects/fastqc/) and sequencing on a NovaSeq 6000 system using a paired-end, 2×150 cycle strategy (Illumina Inc.). The raw data were analysed by NEGEDIA Exome pipeline (v1.0) which involves, alignment to the reference genome (hg38, GCA_000001405.15), removal of duplicate reads and variant calling with Sentieon 202308^92^. Variants were annotated by the Ensembl Variant Effect Predictor (VEP) tool3 (v. 104)^93^. Annotated variants were selected based on: impact on protein High/Moderate, base depth ≧30, alternate depth ≧5 and alt/base depth ≧5%. Selected variants were finally classified with Varsome Clinical (https://eu.clinical.varsome.com/).

### Bulk RNA sequencing and analysis

Total RNA from HPD06 and HPD03 naive hiPSCs was isolated as previously described, and Quant Seq 3’ mRNA-seq Library Prep kit (Lexogen) was used for library construction. Before sequencing library quantification was performed by fluorometer (Qubit) and bioanalyzer (Agilent). Sequencing was performed on NextSeq500 ILLUMINA instruments to produce 5 million reads (75bp SE) for each sample. Transcript quantification was performed from raw reads using Salmon (v1.6.0)^94^ on transcripts defined in Ensembl 106. Gene expression levels were estimated with tximport R package (v1.26.1)^95^.

Batch correction was performed using the ComBat_seq function from the sva R package (v3.50.0; https://bioconductor.org/packages/release/bioc/html/sva.html). Batches have been defined following library preparation: batch 1 for full-length libraries and batch 2 for Quant Seq 3′ mRNA-seq Library Prep kit. CPM on batch corrected counts were computed using the CPM function of the edgeR package (v4.0.12^96^). PCA was performed using the svd r function on log-transformed CPM. All plots except the heatmaps have been done using ggplot2 v3.5.1 (https://ggplot2.tidyverse.org/authors.html#citation). Heatmap was done using the pheatmap R package (v1.0.12). All analyses were performed using R v4.3.2.

Imprinted genes analysis of HPD00 primed and HPD06 and HPD03 naive hiPSCs have been performed by submitting the .fastq raw files to BrewerIX (https://brewerix.bio.unipd.it)^45^, implementing the complete pipeline and evaluating the significant genes.

Total RNA from naive hESCs Shef6 was isolated as previously described. Library preparation with the Quant Seq 3’ mRNA-seq Library Prep kit (Lexogen) and sequencing with the Illumina NextSeq2000 P3 platform was carried out at the VBCF NGS facility, producing 5-10 million reads (50 bp SE). For analysis of the fastq files the Nextflow 23.04.1.5866/nf-core/rnaseq v3.10.1 pipeline was used. This included quality control with fastQC (v0.11.9), pseudo-alignment to the human reference genome hg38 with Salmon (v1.9.0) and alignment with STAR (v2.7.10a). Gene expression levels were estimated with the tximport R package (v1.26.1)^95^. DESeq2 (v1.38.3) was used for further analysis including differential expression analyses. For comparison with already published data^68^, the batch correction was performed using the ComBat_seq function from the sva R package (v3.46.0). For PCAs, counts were transformed with the regularised log transformation (rlog) function (for own time course) or the variance-stabilising transformation (vst) function (integration with Rostovskaya data) from DESeq2. All plots except heatmap have been done using ggplot2 v3.5.1. Heatmaps were done with the pheatmap R package (v1.0.12) using row-wise z-transformed DESeq2-normalized counts. All analyses were performed using R v4.3.2.

### Single-cell RNA sequencing, analysis and comparison with bulk RNA sequencing

For the analyses of the pluripotency exit, the human embryogenesis reference dataset^69^ was derived by the integration of six published human datasets^20,40,75–78^ covering developmental stages from the zygote to the gastrula was kindly provided by the Petropoulos lab. Analysis was performed using Seurat v5.1.0 and the data was subset for embryonic cells covering E5-E14. For the PCA, pseudobulk data was generated with Seurat’s AggregateExpression function and integrated with the bulk RNA-seq data using DESeq2 v1.38.3. Batch correction for the datasets was performed using the ComBat_seq function before applying variance stabilising transformation with the vst function from DESeq2. Heatmaps and boxplots were generated using the Shiny app (http://petropoulos-lanner-labs.clintec.ki.se) from the reference dataset^69^.

For blastoids validation blastoids were dissociated into a single cell suspension by incubation with a mixture of TrypLE™ (Thermo Fisher Scientific A1217701) and Accutase (Merck Sigma Aldrich A6964). The sample was incubated with 0.5 μg of unique hashtag antibody for 20 min on ice. TotalSeq hashtag antibodies (Biolegend) were used to multiplex the samples^97^. The sample was sorted on a BD FACSymphony S6 (BD Biosciences) and then loaded onto a Chromium Next GEM chip (10x Genomics). Further steps of library preparation were performed according to the Chromium Next GEM Single Cell 3’ v3.1 user guide with the addition of the hashtag library for demultiplexing. Combined libraries were sequenced using the NextSeq2000 P2-100 kit (Illumina) with Paired-end sequencing. Initial processing of scRNA-seq data was performed using Cell Ranger (v 6.1.2, 10X Genomics). Dual-indexed RNA and single-indexed hashtag oligo (HTO) libraries were processed in separate instances of cellranger mkfastq, using Illumina’s bcl2fastq (v2.20.0.422). FASTQ files were aligned to the human reference genome (GRCh38, v 2020-A as provided by 10x Genomics) with cellranger multi (expect-cells 16500, min-assignment-confidence 0.9), assigning 4251 cells to the sample of interest. The resulting filtered feature-barcode matrix was loaded into Seurat (v 4.3.0)^98^, excluding features that were detected in less than three cells. Next, RNA data was normalised with the default LogNormalize method, and the HTO assay was normalised with centered log-ratio (CLR) transformation. Based on QC plots, cells with more than 15% mitochondrial counts or less than 7000 UMIs (nCount_RNA) were removed, retaining 3260 high-quality cells. These were subjected to standard Seurat processing using mostly default parameters unless indicated: FindVariableFeatures, ScaleData, RunPCA, RunUMAP (dims=1:15), FindNeighbors (dims=1:15), FindClusters (resolution=0.5). To characterise the cells’ cycling status, the function CellCycleScoring was run using the built-in lists of S and G2M phase markers (cc.genes.updated.2019) derived from^99^. Cells separated by cell cycle phase on the UMAP and this source of heterogeneity was then regressed out using ScaleData (vars.to.regress = c(“S.Score”, “G2M.Score”), features = rownames(object)), followed by RunPCA, RunUMAP(dims=1:15), FindNeighbors(dims=1:15), and FindClusters(resolution=0.5), resulting in 5 clusters. The raw counts of the quality-filtered matrix was uploaded to the Early Embryogenesis Projection Tool (v2.1.1) (https://petropoulos-lanner-labs.clintec.ki.se/shinys/app/ShinyEmbryoProjP, accessed on 2024-09-26) to project the cells on an integrated reference UMAP of human embryo development and get annotation of predicted cell identities^69^.

### Statistics and reproducibility

For each dataset, sample size *n* refers to the number of biological or technical replicates, shown as dots and stated in the figure legends. All error bars indicate the standard error of the mean (SEM). All RT-qPCR experiments were performed in three technical replicates.

### Ethics statement

Our research complies with all relevant ethical regulations including ISSCR guidelines. HPD06 and HPD03 naive hiPSCs used in G.M.’s Laboratory were checked by the European ethics committee and registered in the human pluripotent stem cells registry (link: https://hpscreg.eu/ cell-line/UNIPDi004-B). Experiments with hPSCs and blastoids in J.J.Z’s Laboratory were approved by the Scientific Ethics Committee for Hovedstaden (H-21043866/94634 and H-24048289). Shef6 line use is with the agreement of the Steering Committee of the UK Stem Cell Bank (SCSC16-09). The WiCell line H9 (WA09) was used under the agreements 23-W0460 (JJZ) and 21-W0002 (ML). Work with human embryonic and induced pluripotent stem cells including blastoids in V.P. Laboratory was approved by the UZ/KU Leuven ethics committee (S64962, S66595 and S68981). Work with mouse embryonic fibroblast was approved by the UZ/KU Leuven ethics committee (P170/2019). The Austrian Academy of Sciences (the local ethical body) has given N.R.’s Laboratory a license to perform blastoid experiments, following expert legal advice that concluded these are not in conflict with Austrian laws. This license is in the shape of a statement of the Commission for Science Ethics of the Austrian Academy of Sciences concerning the project ‘Modeling human early development using stem cells’. There is no approval number on that document. This license conforms with the ethical standards suggested by the International Society for Stem Cell Research (ISSCR). This work did not exceed a developmental stage normally associated with 14 consecutive days in culture after fertilization nor did it entail any implantation *in vivo*.

### Inclusion & ethics statement

All collaborators of this study have fulfilled the criteria for authorship required by Nature Portfolio journals and have been included as authors, as their participation was essential for the design and implementation of the study. Roles and responsibilities were agreed among collaborators ahead of the research. This research was not severely restricted or prohibited in the setting of the researchers and does not result in stigmatisation, incrimination, discrimination or personal risk to participants.

## Data availability

Bulk and single-cell RNA-seq data for this study have been deposited in the Gene Expression Omnibus (GEO) database under the accession code GSE284370. We also included available RNA-Seq data for naive and primed hPSCs, hTSCs and fibroblasts from GSE110377^30^, GSE63577^100^, GSE93226^101^, GSE75868^43^, GSE184562^72^, GSE138688^34^, GSE178162^102^, GSE135695^103^, PRJNA397941^27^, GSE133630^104^, GSE73211^105^, PRJEB7132^24^, PRJEB12748^26^; for capacitation from GSE123055^68^; for embryonic development from the human embryogenesis reference dataset^69^, comprehensive of GSE36552^20^, E-MTAB-3929^75^, GSE136447^76^, E-MTAB-9388^78^, GSE171820^40^, PRJEB30442^77^. Proteomics data have been deposited in the PRIDE Proteomics database under the accession code PXD059820.

## Acknowledgements

We thank the KU Leuven FACS Core team for supporting flow cytometry experiments, Alejandra Purcell from Gorgas Institute in FACS analysis, and the VIB BioImaging Core Leuven for their contribution to imaging performed in V.P. lab. We thank the Vienna Biocenter Core Facilities (VBCF) for support in NGS analysis. J.J.Z’s team thanks Antar Drews and Sandra Bages Arnal for their input, the reNEW platforms for technical expertise, support and use of equipment in particular: H. Wollmann, M. Michaut, J. Bulkescher, G. Dela Cruz and A. Kalvisa.

G.M.’s Laboratory is supported by grants from: Giovanni Armenise–Harvard Foundation, Telethon Foundation (GJC21157), European Research Council Starting Grant (716910 - MetEpiStem), Italian national project ‘PRIN 2022’ (2022RA8E3T) and La Caixa Foundation (LCF/PR/HR24/52440015). G.R. is supported by a HORIZON MSCA Postdoctoral Fellowship (101108873 - PLURImet). M.L.’s Laboratory is supported by grants from the Austrian Science Fund (FWF; P35637, I5958). M.O. is a member of the FWF-funded doctoral programme ‘Signalling Molecules in Cellular Homeostasis’ (SMICH; W1261), and M.L. is a faculty member and speaker of SMICH and a ‘Wiener Wissenschafts-Forschungs- und Technologiefonds (WWTF)’ Vienna Research Group Leader (VRG14-006). V.P.’s Laboratory is supported by the Research Foundation-Flanders (FWO grants G0C9320N and G0B4420N to V.P.), KU Leuven Research Fund (C1 grant C14/21/119 to V.P.), and Pandarome project 40007487 (G0I7822N) (funded by the FWO and F.R.S.-FNRS) under the Excellence of Science (EOS) program. M.A.S. is supported by Gorgas Memorial Institute for Health Studies and Fundación Sus Buenos Vecinos in Panama. S.S.F.A.v.K. (11I1523N) and T.X.A.P. (11N3122N) are supported by FWO PhD fellowships. J.J.Z.’s Laboratory is supported by grants from: Novo Nordisk Fonden (NNF21CC0073729), Lundbeckfonden (R345-2020-14979), Danmarks Frie Forskningsfond (0169-00031B) and the European Research Council Starting Grant (101077271 - ChroMeta). N.R.’s Laboratory is supported by the European Research Council (ERC) under the European Union’s Horizon 2020 research and innovation programme (ERC-Co grant agreement no. 101002317, “BLASTOID: a discovery platform for early human embryogenesis”).

D.C. is supported by Fondazione Telethon Core Grant, Italian Ministry of Health (Piano Operativo Salute Traiettoria 3, T3-AN-09, “Genomed”; Ricerca Finalizzata 2021, “genOMICA”; MCNT2 2023, “EUCARDIS”), Italian Ministry of University and Research and European Union (Next Generation EU - MUR-PRIN-2022, Project PNC 0000001 D3 4 Health). P.G. is supported by Fondazione Telethon Core Grant, AIRC (MFAG 2020), PNRR (PNRR-MR1-2022-12376821), PRIN-2022-PNRR (P2022JLNZ7), PRIN (20224FL9T5). P.M. is supported by the Italian national project ‘PRIN 2022’ (2022LJZRBY).

## Author contributions

G.R. performed experiments and analysed data (characterisation, embryonic and TSCs differentiation, bulk RNA-Seq, proteomics), supervised the study, prepared figures and wrote the manuscript; I.Z., performed experiments (characterisation) and edited the manuscript; A.D. performed experiments (characterisation); G.P. and N.M. performed experiments (characterisation, embryonic and TSC differentiation); P.M. analysed bulk RNA-Seq for the comparison with published datasets; C.C. and D.C. generated and analysed Exome-Seq data; P.G. performed proteomics experiments; G.M. conceived and supervised the study, secured funding and wrote the manuscript. M.O. performed experiments and analysed NGS data (embryonic differentiation), wrote the manuscript and prepared figures; M.L. supervised the study, secured funding and wrote the manuscript. A.O. and E.v.G. performed experiments (blastoids); M.A.S., S.S.F.A.v.K., T.X.A.P., and S.K. performed experiments (characterisation, TSC differentiation); N.R. and V.P. supervised the study, edited the manuscript and secured funding. I.S.B. performed experiments (blastoids), analysed the data, prepared figures and edited the manuscript; A.W. analysed and curated scRNAseq data; J.J.Z. supervised the study, secured funding, prepared figures and wrote the manuscript.

## Supplementary Material

**Supplementary Table 1.**
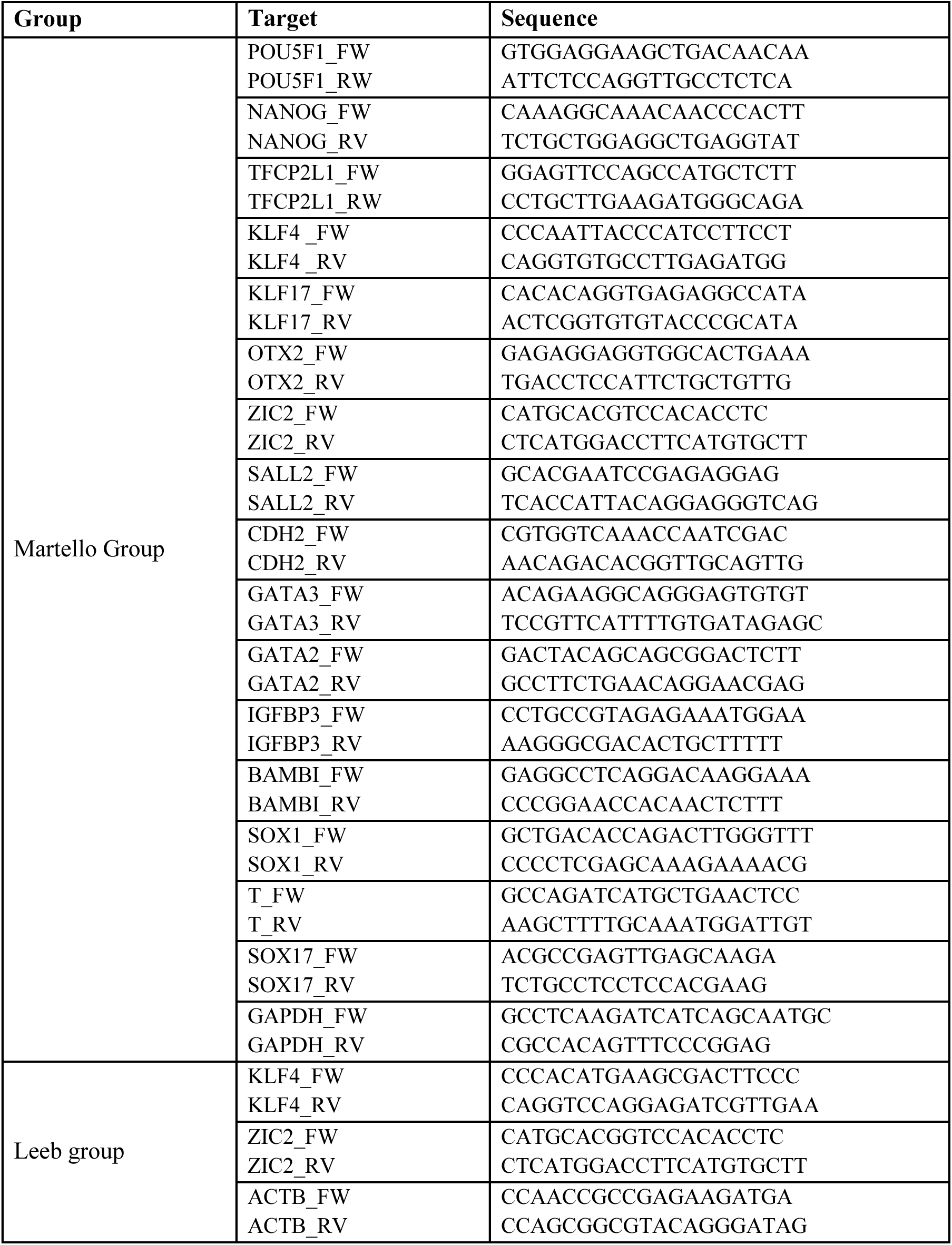
Primers used in this study.

**Supplementary Table 2.** Primary antibodies used in this study.

| Group | Target | Conjugate | Supplier | Catalog ID | IF dil. |
| --- | --- | --- | --- | --- | --- |
| Martello Group | KLF17 | - | Atlas Antibodies | HPA024629 | 1:250 |
|  | OCT4 (C-10) | - | Santa Cruz | sc-5279 | 1:300 |
|  | NANOG | - | Cell Signaling | D73G4 | 1:100 |
|  | TFCP2L1 | - | R&D Systems | AF5726 | 1:250 |
|  | GATA3 | - | R&D Systems | AF2605 | 1:100 |
| Leeb Group | KLF17 | - | Atlas Antibodies | HPA024629 | 1:333 |
|  | OCT4 (C-10) | - | Santa Cruz | sc5279 | 1:50 |
|  | SUSD2 (W5C5) | APC | BioLegend | 327408 | 1:100 |
| Pasque Group | KLF17 | - | Abcam | HPA024629 | 1:300 |
|  | OCT4 (C-10) | - | Santa Cruz | sc-5279 | 1:100 |
|  | NANOG | - | BD Bioscience | 560482 | 1:50 |
|  | GATA3 |  | R&D System | AF2605 | 1:100 |
|  | SOX17 | - | R&D System | AF1924 | 1:100 |
|  | SUSD2 | PE | Biolegend | 327406 | 1:50 |
|  | GATA2 | - | Sigma-Aldrich | WH0002624M1 | 1:100 |
|  | NANOG | - | Abcam | ab21624 | 1:100 |
| Zylicz Group | NANOG | - | Abcam | ab173368 | 1:250 |
|  | GATA3 | - | Abcam | ab199428 | 1:250 |
|  | GATA4 | - | eBioscience | 14-9980-80 | 1:250 |
| Rivron Lab | OCT4 (C-10) | - | Santa Cruz | sc-5279 | 1:200 |
|  | CDX2 | - | Abcam | ab76541 | 1:1000 |
|  | GATA4 | - | Invitrogen | 14-9980-82 | 1:400 |

**SUPPLEMENTARY FIGURE 1.**
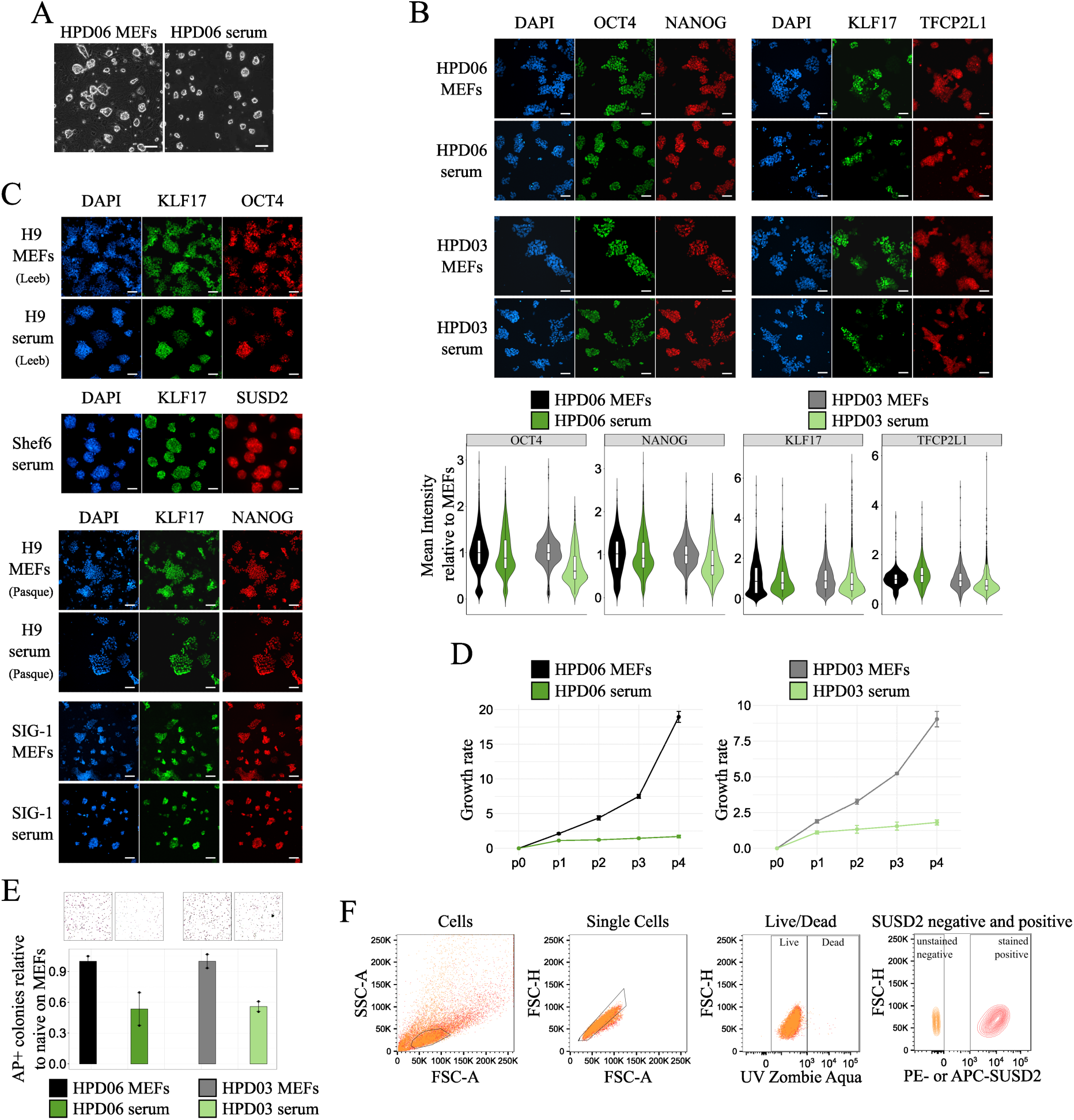
**A** Morphologies of HPD06 naive hiPSC line cultured on MEFs or serum coating at the 4th passage when plated at a low density. Scale bars: 100 μm. Representative images of two independent experiments are shown. **B** Top: Immunostaining for general pluripotency (OCT4 and NANOG) and naive (KLF17 and TFCP2L1) markers of HPD06 and HPD03 naive hiPSCs cultured on MEFs or serum coating at the 4th passage. Complete view of Figure 1B. Scale bars: 100 μM. Representative images of two independent experiments are shown. Bottom: Mean fluorescence intensity quantification for general pluripotency (OCT4 and NANOG) and naive (KLF17 and TFCP2L1) markers of HPD06 and HPD03 naive hiPSCs cultured on MEFs or serum coating at the 4th passage. At least 700 nuclei from five randomly selected fields from two independent experiments were analysed for each condition. The box plot indicates the 25th, 50th and 75th percentiles. **C** Immunostaining for general pluripotency (OCT4 and NANOG) and naive (KLF17 and SUSD2) markers of H9 and Shef6 naive hESCs, and SIG-1 hiPSCs stably cultured on MEFs or serum coating. Complete view of Figure 1B. Scale bars: 100 μM. Representative images of two independent experiments are shown. **D** Growth rate of HPD06 and HPD03 naive hiPSCs cultured on MEFs or serum coating when plated at a low density over the first 4 passages of the conversion. Bars indicate the mean ± SEM of n = 2 independent experiments shown as dots. **E** Top: Representative AP staining images after clonal assay of HPD06 and HPD03 naive hiPSCs cultured on MEFs or serum coating when plated at a low density at the 4th passage. Bottom: Quantification of the relative number of AP-positive pluripotent colonies counted per well. Bars indicate the mean ± SEM of n = 3 independent experiments shown as dots. **F** Representative gating strategy to evaluate SUSD2 positivity in naive hPSCs by flow cytometry. Selected sub-populations are shown from left to right. First, the cell population was distinguished from cell debris (left panel). Singlets were chosen from the cell population and live cells among singlets were selected, followed by the gating of SUSD2-positive cells using unstained negative control (right panel). This gating strategy corresponds to Figure 1G and Supplementary Figure 3F.

**SUPPLEMENTARY FIGURE 2.**
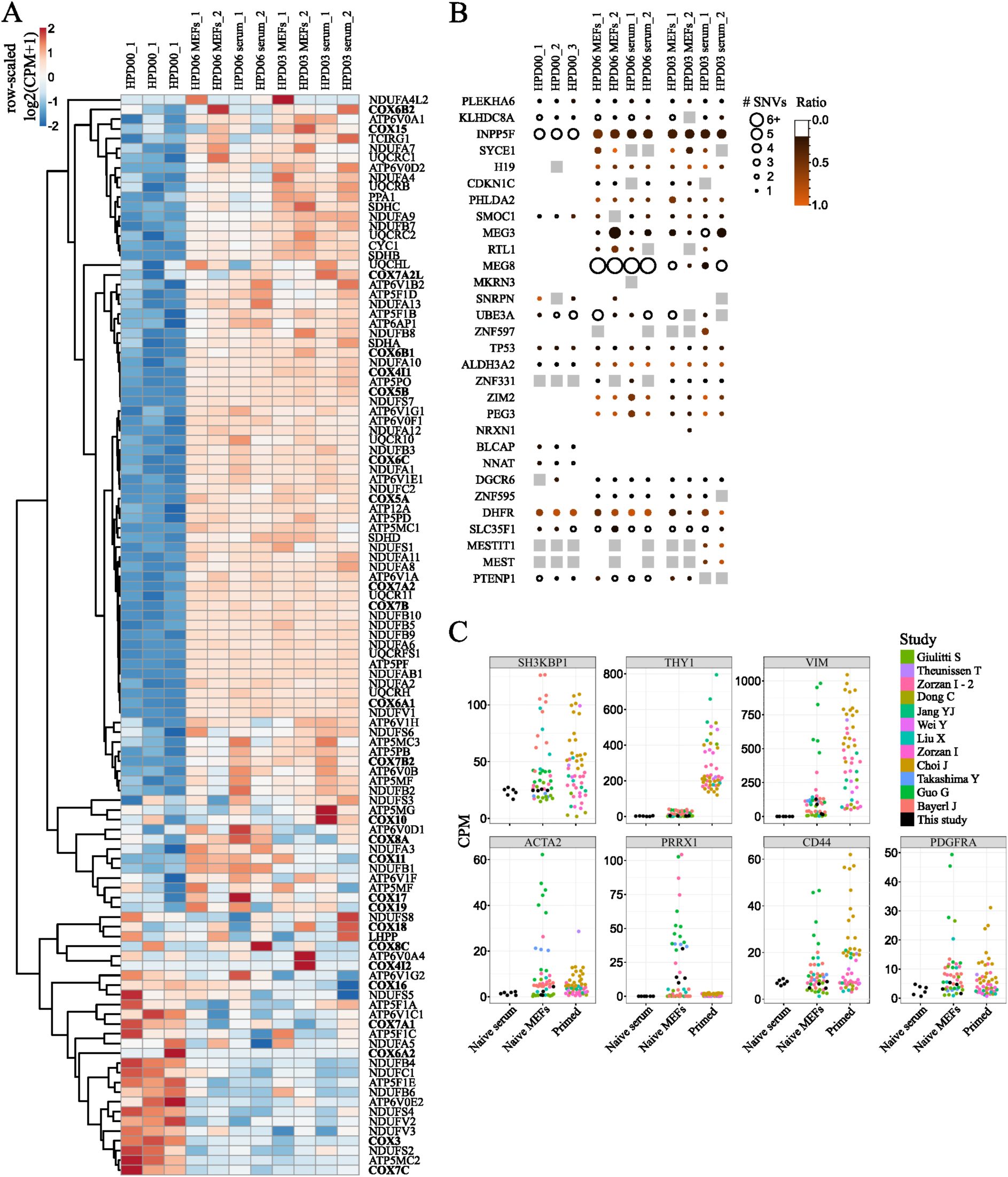
**A** Heatmap of oxidative phosphorylation genes from the KEGG PATHWAY Database (https://www.genome.jp/kegg/pathway.html) and COX genes^24^ in HPD00 primed hiPSCs and HPD06 and HPD03 naive hiPSCs stably cultured on MEFs or serum coating. **B** BrewerIX gene summary panel results on RNA-Seq data from HPD00 primed hiPSCs and HPD06 and HPD03 naive hiPSCs stably cultured on MEFs or serum coating. Empty dots indicate detected genes with no evidence of biallelic expression, grey squares indicate genes detected but not reaching the thresholds, and the absence of any symbol indicates that the gene was not detected. **C** Absolute expression (CPM) of fibroblasts markers from Figure 2D in naive hPSC lines (HPD06 and HPD03 hiPSCs, and Shef6 hESCs) stably cultured on MEFs or serum coating and in published naive hPSCs on MEFs and primed hPSCs.

**SUPPLEMENTARY FIGURE 3.**
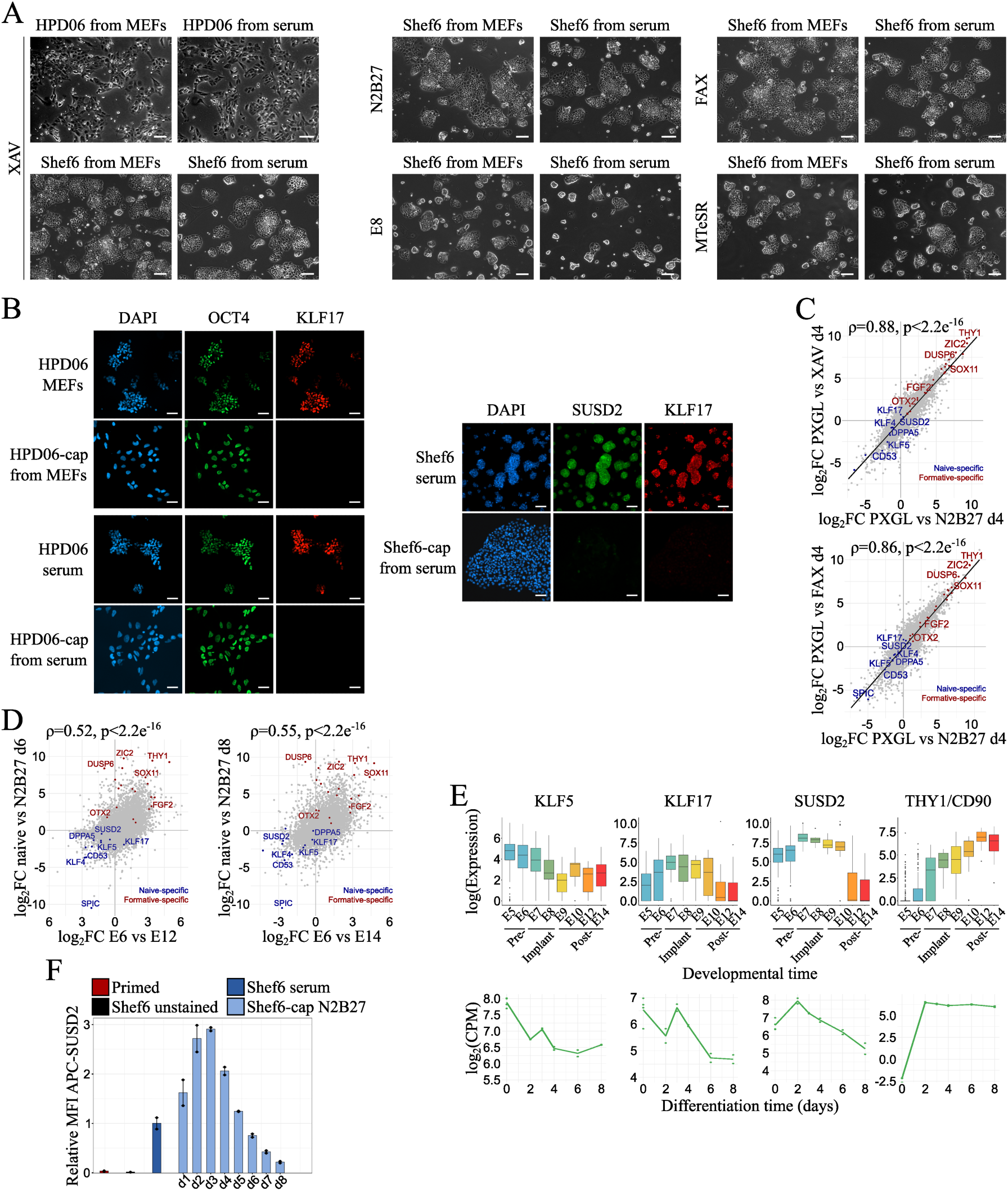
**A** Morphologies of HPD06 hiPSCs and Shef6 hESCs capacitated from naive lines stably cultured on MEFs or serum coating under different media compositions. Scale bars: 100 μm. Representative images of two independent experiments are shown. **B** Immunostaining for general (OCT4) and naive (KLF17 and SUSD2) pluripotency markers of HPD06 hiPSCs and Shef6 hESCs capacitated from naive lines stably cultured on MEFs or serum coating. Scale bars: 100 μM. Representative images of two independent experiments are shown. **C** Scatter Plots showing log2 fold changes (log2FC) of Shef6 naive hESCs stably cultured on serum coating differentiated for 4 days in unsupplemented N2B27 (n = 4 from two independent experiments), N2B27 supplemented with XAV (n = 2 biological replicates), or N2B27 supplemented with XAV, FGF2, and Activin A (FAX, n = 2 biological replicates) compared to the naive state (n = 6 from two independent experiments). Data was filtered for the 9376 DEGs (|log2FC| > 1, padj < 0.05) identified in any comparison with the naive state. Spearman correlation (ρ, n = 9376) and approximated p-values are overlaid, with a linear regression line (for visualization) fitted. Selected naive- and primed-specific markers are labelled. **D** Scatter Plots showing log2 fold changes (log2FC) of Shef6 naive hESCs stably cultured on serum coating in the naive state (n = 4 from two independent experiments) versus differentiation for 6 (n = 2 biological replicates) or 8 (n = 2 biological replicates) days in N2B27, compared to corresponding developmental times (E6 vs. E12 or E14) from the human embryonic reference dataset (subset for embryonic lineages only)^75^. Data was filtered for the 8508 DEGs (|log2FC| > 1, padj < 0.05) identified in any comparison between naive hESCs in PXGL and any differentiation timepoint in N2B27, as well as between E5 or E6 and any later embryonic day until E14. Spearman correlation (ρ, n = 8508) and approximated p-values are overlaid, with selected naive- and primed-specific genes labelled. **E** Gene expression of naive (*KLF5*, *KLF17* and *SUSD2*), and primed (*THY1*) pluripotency markers in the human embryonic reference dataset (subset for embryonic lineages only)^75^ and for Shef6 naive hESCs stably cultured on serum coating differentiated for 6-8 days in N2B27 measured by RNA-seq. Lines indicate inferred trends based on mean values, with biological replicates (n = 4 from 2 independent experiments for d0, n = 2 biological replicates for any other time point) shown as dots. **F** Expression of the naive-specific surface marker SUSD2 in Shef6 naive hESCs stably cultured on serum coating during capacitation for 8 days in N2B27, measured by flow cytometry using an APC-conjugated anti-SUSD2 antibody. The y-axis represents the relative median fluorescence intensity (MFI) normalised to the naive state. Bars represent the mean ± SEM of n = 2 biological replicate shown as dots.

**SUPPLEMENTARY FIGURE 4.**
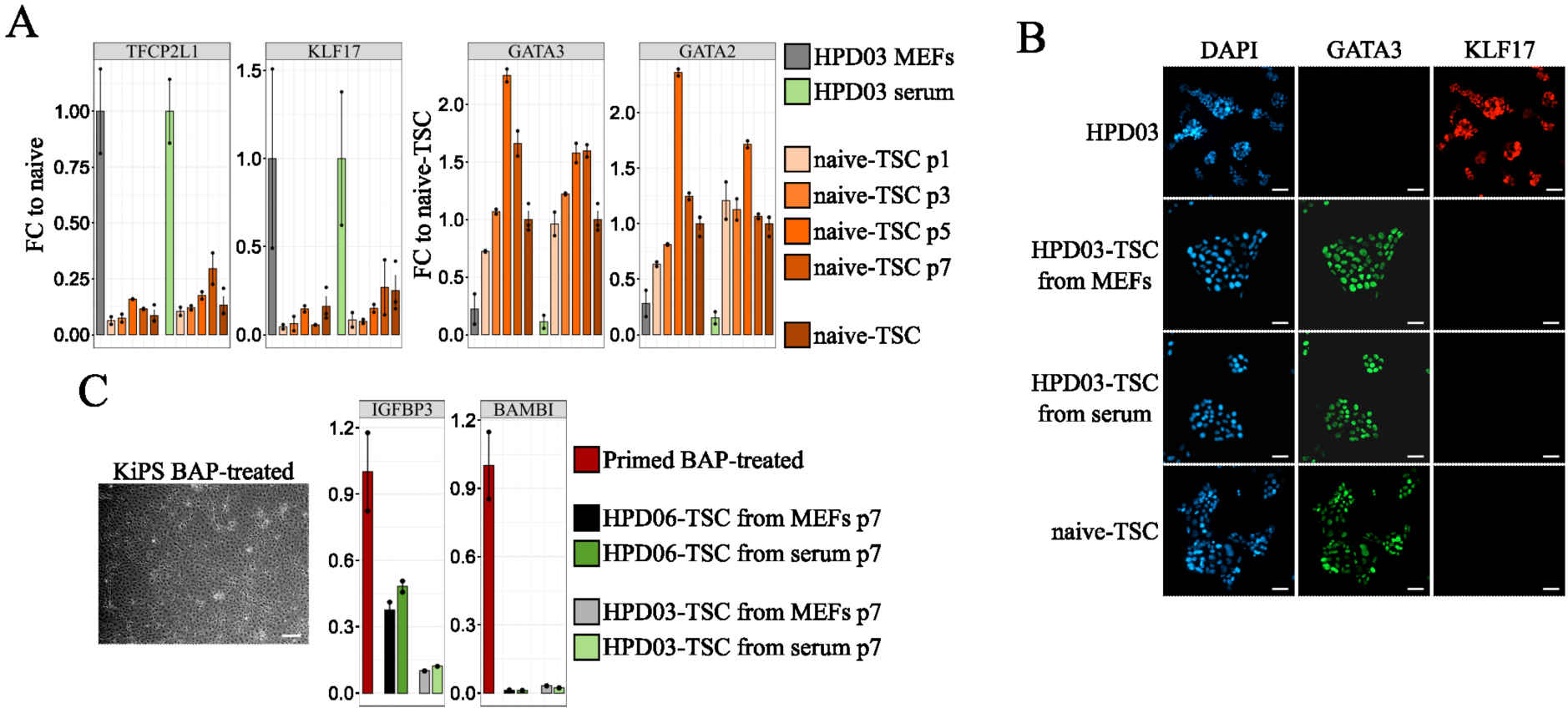
**A** Gene expression analysis by RT-qPCR of naive pluripotency (*TFCP2L1* and *KLF17*) and TSCs (*GATA3* and *GATA2*) markers in TSCs derived from HPD03 naive hiPSCs stably cultured on MEFs or serum coating. Bars indicate the mean ± SEM of technical replicates shown as dots. Biological replicates include n = 3 independent experiments for naive hPSCs and naive-TSCs, and n = 2 independent experiments for the differentiation time points. **B** Immunostaining for TSCs (GATA3) and naive pluripotency (KLF17) markers in TSCs derived from HPD03 naive hiPSCs stably cultured on MEFs or serum coating. Scale bars: 100 μM. Representative images of two independent experiments are shown. **C** Left: Morphology of KiPS primed hiPSCs treated with BAP medium for 4 days. Scale bar: 100 μm. Right: Gene expression analysis by RT-qPCR of amnion markers (*IGFBP3* and *BAMBI*) in TSCs derived from HPD06 and HPD03 naive hiPSCs stably cultured on MEFs or serum coating. Bars indicate the mean ± SEM of n = 2 biological replicates shown as dots.

**SUPPLEMENTARY FIGURE 5.**
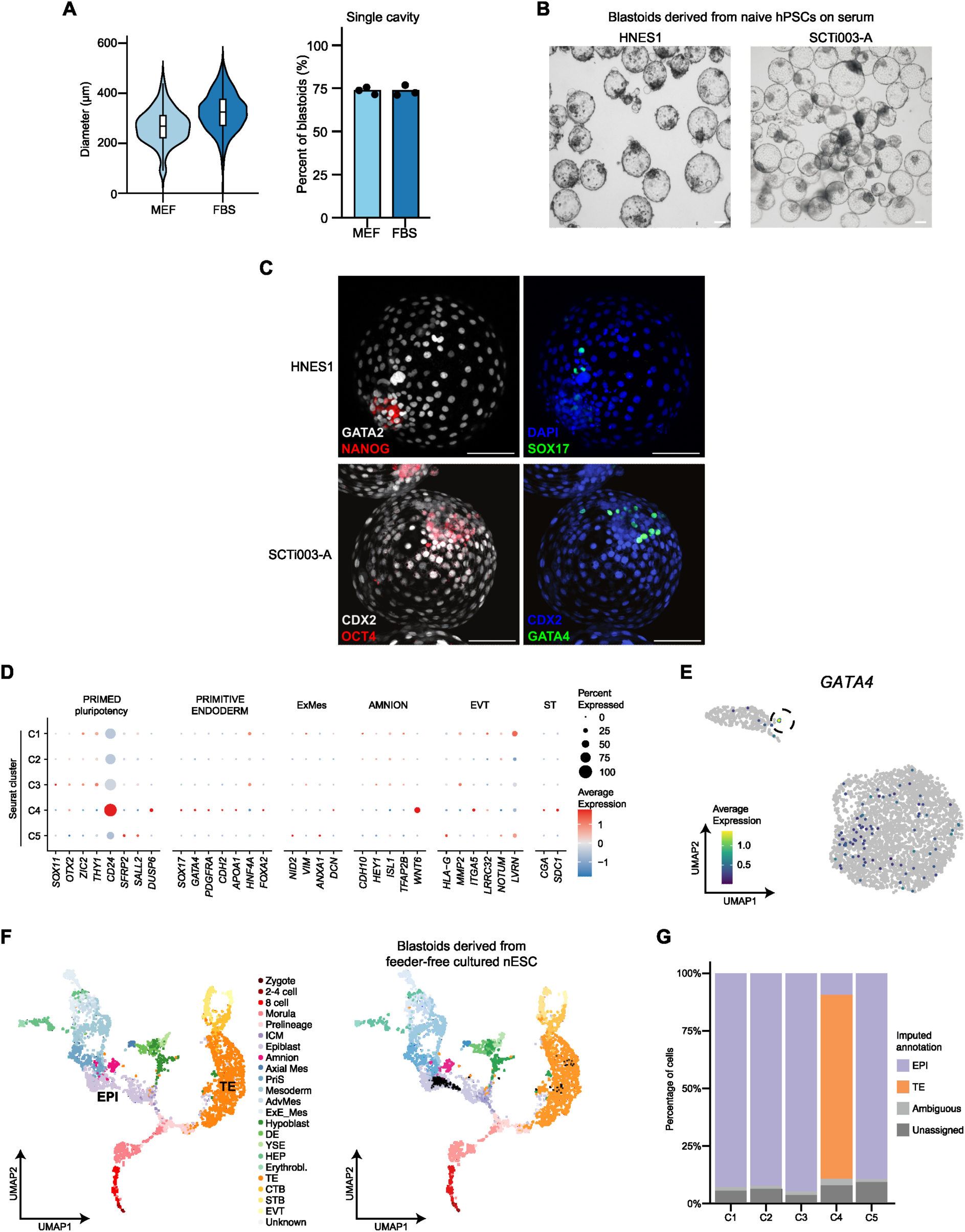
**A** Quantification of blastoid diameter (left) and blastoids with a single cavity (right) in structures formed from naive H9 hESCs cultured on MEFs or serum coating. Left: data is presented as violin plots with the median, 25, 75 percentiles (± min-max) of n = 407 (MEFs) and n = 806 (serum coating) across 3 biological replicates. Right: Data shows the mean percentages and data points of 3 biological replicates. **B** Brightfield images of blastoids induced from naive HNES1 hESC and naive SCTi003-A hiPSCs cultured on serum coating. Scale bar: 100 μm. **C** Immunostaining of blastoids derived from naive HNES1 hESC and naive SCTi003-A hiPSCs cultured on serum coating. Blastoids were stained for TE (GATA2 or CDX2), PrE (GATA4 or SOX17) and EPI (NANOG and OCT4) markers. Shown is the maximum projection. Scale bars: 100 μm. **D** Expression of selected lineage-specific marker genes across Seurat clusters from Figure 6C. The size of the dots represents the proportion of cells in the indicated group expressing the given gene and colour encodes the scaled average expression. **E** UMAP plot from scRNA-seq analysis as in Figure 6C. Cells are coloured according to *GATA4* expression (n = 3260). **F** Left: UMAP of human pre-implantation and post-implantation embryos with annotation for scRNA-seq data integration from^69^ with cells coloured by cell type. Right: Projection of in vivo reference data with in vitro d5 blastoids from naive H9 hESC cultured on serum coating. Black dots show neighbourhoods of in vitro generated cells projected onto a reference UMAP. **G** Imputed cell annotation across different Seurat clusters as in Figure 6F. Unassigned and ambiguous labels refer to cells with either none or with more than two imputed stages respectively.

